# iPAR: a new reporter for eukaryotic cytoplasmic protein aggregation

**DOI:** 10.1101/2024.01.29.577793

**Authors:** Sarah Lecinski, Jamieson A.L. Howard, Chris MacDonald, Mark C. Leake

## Abstract

**Background:** Cells employ myriad regulatory mechanisms to maintain protein homeostasis, termed proteostasis, to ensure correct cellular function. Dysregulation of proteostasis, which is often induced by physiological stress and ageing, often results in Protein Aggregation in cells. These aggregated structures can perturb normal physiological function, compromising cell integrity and viability, a prime example being early onset of several neurodegenerative diseases. Understanding aggregate dynamics *in vivo* is therefore of strong interest for biomedicine and pharmacology. However, factors involved in formation, distribution and clearance of intracellular aggregates are not fully understood.

**Methods:** Here, we report an improved methodology for production of fluorescent aggregates in model budding yeast which can be detected, tracked and quantified using fluorescence microscopy in live cells. This new openly-available technology, iPAR (inducible Protein Aggregation Reporter), involves monomeric fluorescent protein reporters fused to a ΔssCPY* aggregation biomarker, with expression controlled under the copper-regulated *CUP1* promoter.

**Results:** Monomeric tags overcome challenges associated with non-physiological reporter aggregation, whilst *CUP1* provides more precise control of protein production. We show that iPAR and the associated bioimaging methodology enables quantitative study of cytoplasmic aggregate kinetics and inheritance features *in vivo*. We demonstrate that iPAR can be used with traditional epifluorescence and confocal microscopy as well as single-molecule precise Slimfield millisecond microscopy. Our results indicate that cytoplasmic aggregates are mobile and contain a broad range of number of iPAR molecules, from tens to several hundred per aggregate, whose mean value increases with extracellular hyperosmotic stress.

**Discussion:** Time lapse imaging shows that although larger iPAR aggregates associate with nuclear and vacuolar compartments, and for the first time we show directly that these proteotoxic accumulations are not inherited by daughter cells, unlike nuclei and vacuoles. If suitably adapted, iPAR offers new potential for studying diseases relating to protein oligomerization processes in other model cellular systems.

## 1. Introduction

Accumulation of misfolded protein aggregates is triggered by environmental stress conditions, which in turn compromise cell function. However, cells have evolved to respond to these changes to maintain metabolic function and ensure survival. In eukaryotic cells, systems such as the temporal protein quality control (PQC) sustain the proteome and actively contribute to the detection of misfolded proteins (1, 2), promoting their refolding mediated by chaperone proteins (2, 3). The degradation of damaged proteins is actively mediated by the ubiquitin-proteasome system (UPS) (4, 5) but not all proteins are recognised this way, and other selective processes exist to degrade proteins, such as the autophagy pathway (6). Generally, these systems require acute control of the temporal and spatial dynamics of subcellular components for quality control *in vivo* to prevent or clear aggregates and maintain proteomic homeostasis (2, 3, 5).

When quality control responses and processes fail, misfolded proteins accumulate in the intracellular environment with a heterogeneous size distribution of aggregates (7, 8), consistent with diffusion-nucleation mechanisms of formation (9). This distribution of protein aggregates is harmful to the cell (10, 11), with endogenous protein aggregation effectively depleted from the cellular environment. Further toxicity is mediated by aggregation through perturbation of other functional proteins present in the crowded intracellular environment (12, 13). Ultimately, this can lead to pathogenic phenotypes (14, 15). Many neurodegenerative diseases (e.g. Parkinson’s and Alzheimer’s) are associated with a process which involves aggregation of amyloid resulting in packed beta-sheet structures and fibres (16–18), due in part to amyloid-β oligomerization (19). Other diseases such as cataracts (20) and Huntington’s disease (21) result from the formation of amorphous aggregates (22, 23). Understanding the formation of such proteotoxic factors is crucial to elucidating underlying mechanisms associated with cellular malfunction and toxicity. Insight into the associated *in vivo* dynamics of these factors can also contribute to the development of new therapeutic methods.

Budding yeast, *Saccharomyces cerevisiae*, has been used to investigate several important processes affecting intracellular organisation which are highly conserved across all eukaryotes, including key survival mechanisms (24, 25), essential metabolic pathways such as DNA replication (26, 27), transcription (28, 29), membrane trafficking (30–33), and PQC machinery for aggregate detection and clearance (3, 34, 35). Considering its excellent genetic tractability, and ease of cell culturing and optical imaging, we used *S. cerevisiae* as a eukaryotic cellular model to investigate intracellular dynamics of aggregation. Various markers for aggregation use key conserved proteins present in yeast. Chaperone proteins are a good example of this; considered a first response against misfolded proteins, they are recruited at the site of misfolded proteins or aggregates to promote re-folding or initiate degradation pathways if necessary (36, 37). Current approaches to analysing and quantifying protein aggregates include optical microscopy with use of fluorescent biomarkers of aggregation, typically using chaperone proteins as reporters (e.g. Hsp70, Hsp40, Hsp104) (20, 38–40). Additionally, variants prone to form aggregates have been fluorescently tagged, such as the thermosensitive mutant of Ubc9 (41) derived from a SUMO-conjugating enzyme and unable to properly fold in yeast cells (42).

Another common marker for aggregation used in *S. cerevisiae* is the engineered reporter ΔssCPY*, a misfolded version of the vacuolar enzyme carboxypeptidase Y (CPY), which is prone to form aggregates and mis-localises to the cytoplasm (43, 44). This variant, derived from the native CPY (45, 46), carries a single amino acid mutation with a glycine to arginine substitution at residue position 255 (G255R) (44, 47) (Figure 1). This mutation (labelled CPY*) is responsible for its misfolding, and when combined with an N-terminal truncated signal peptide (Δss) results in aberrant localisation of this misfolded protein to the cytoplasm. Tagging of ΔssCPY* with enhanced GFP (EGFP) has been used as a model to uncover PQC (48–50) and protein sorting dynamics (40, 51), cellular perturbations and protein aggregation kinetics in stressed cells (52, 53). Studies have revealed that protein aggregate interactions and localisation *in vivo* have a crucial role in establishing toxicity (53).

**Figure 1:**
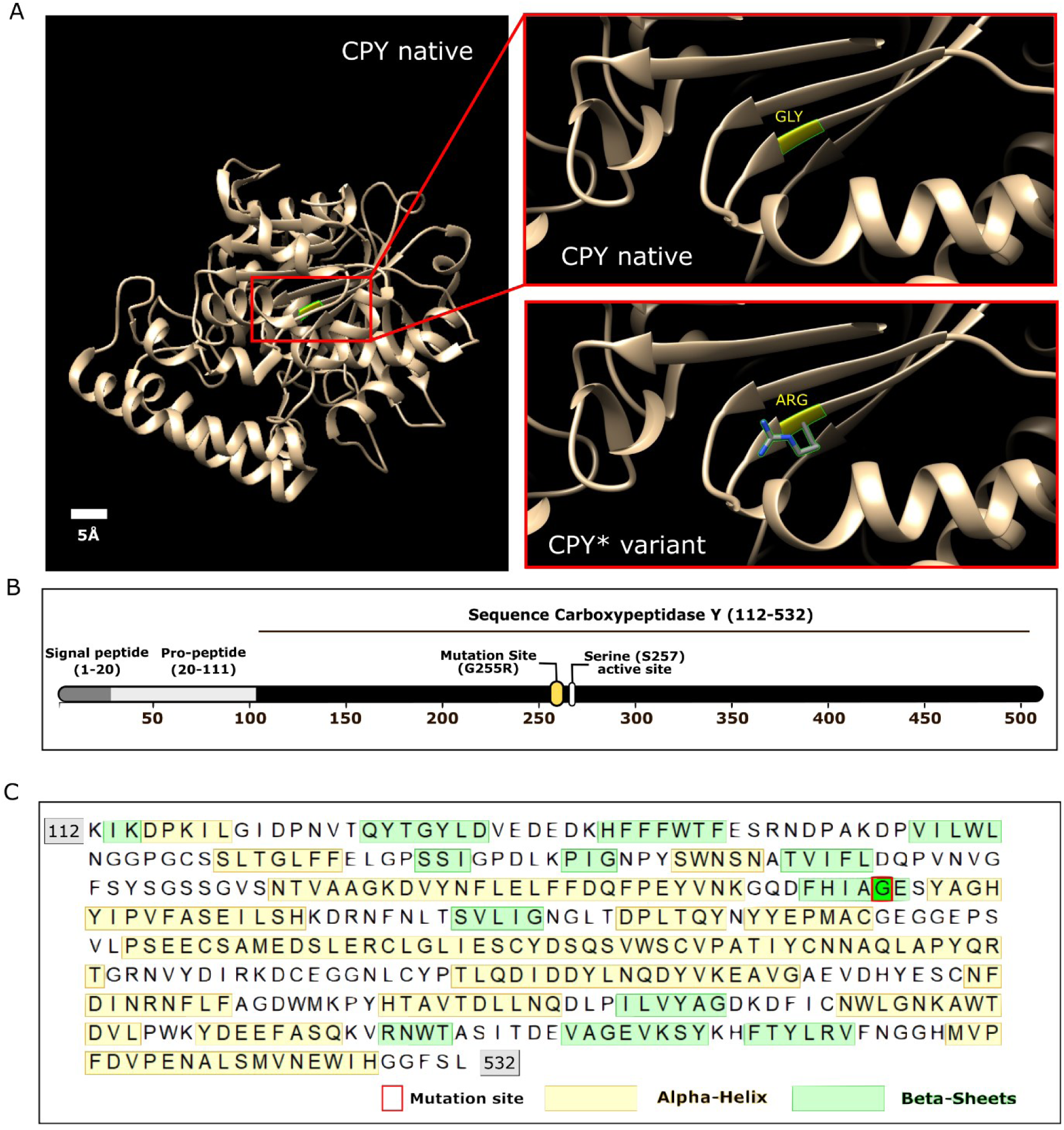
Modifications of CPY to enable its use as a reporter of cytoplasmic protein aggregation. A) Left; a 3D model of the native CPY structure. Right; zoom-in of the mutated region, showing Glycine residue 255 in the native protein and the arginine substitution in the misfolded CPY* variant. Both amino acids are indicated in yellow. The 3D crystal structure of CPY (PDB ID: 1WPX) was visualised using Chimera software. B) CPY sequence showing G255R mutation site near the S257 active site, responsible for the protein unfolding and aggregative behaviour. C) Sequence for CPY, mutation site and native secondary structures. A red rectangle indicates the position of the mutation site (G255), alpha-helix regions in the native protein are shown in yellow and beta-sheets regions are displayed in green.

The ΔssCPY* aggregation reporter is typically expressed from the endogenous *PRC1* promotor, which is problematic as this gene is metabolically regulated, for example being upregulated under certain stress conditions, such as nutrient starvation (54, 55). As protein aggregation correlates with cellular abundance of proteins and local protein concentrations, and is often assessed under stress conditions, there are challenges in disentangling phenotypes which are associated with metabolic-dependent expression and protein aggregation in such experiments. Furthermore, EGFP, and indeed several other fluorescent protein tags, has the capacity to dimerize (56–58), which can also potentially introduce challenging artifacts when assessing the aggregation of tagged molecules.

To address the limitations of existing aggregation biomarkers, we present newly developed versions of ΔssCPY* as reporters for cytoplasmic protein aggregation that are tagged with monomeric fluorescent proteins and are expressed under the control of an inducible promoter. This new class of novel reagent, which we denote as an inducible Protein Aggregation Reporter (iPAR) is part of a useful methodology when used in conjunction with a range of fluorescence microscopy modalities to study several mechanistic aspects of stress-induced protein aggregation in cells.

For iPAR, we replaced the metabolically regulated endogenous promoter (*PRC1*) used to express ΔssCPY* as reporter with the copper inducible promoter (*CUP1)*. The fluorescent fusion tag EGFP was additionally mutated to a monomeric version (mEGFP), which uses electrostatic repulsion to inhibit interactions between pairs of fluorescent protein molecules thereby minimising tag-induced oligomerisation effects. To enhance the utility of this new aggregation reporter with newer developed fluorescent proteins that have brighter fluorescence signal properties and faster maturation times than EGFP (59), we also constructed two variants of iPAR by swapping the mEGFP fusion tag with a brighter green monomeric fluorescent protein mNeonGreen as well as with the red fluorescent protein mScarlet-I. To increase the wider utility of this methodology for researchers, we further made these probes available for expression in budding yeast by creating plasmids of all three iPARs with both *URA3* and *LEU2* selection markers. Using these reagents with newly designed image analysis techniques, we were able to quantify the induced protein aggregation following hyperosmotic and elevated temperature cell stresses, and also to assess the capacity for mother cells to retain protein aggregates during the process of asymmetric cell division, during which other cellular organelles such as the nucleus and lytic vacuole are inherited in budding daughter cells. We further visualise iPAR *in vivo* using Slimfield microscopy, a rapid fluorescence imaging modality which can detect single fluorescent dye molecules including fluorescent proteins in the cytoplasm of a range of organisms including single bacteria (9, 60–67), yeast (68–70), algae (71, 72) and mammalian (73, 74) cells, as well as animal (75) and plant (76) tissue, with below millisecond sampling capability (77). Analysis of iPAR aggregate Slimfield tracks indicate that aggregates are mobile in the vacuolar and nuclear compartments and possess between a few tens and a few hundred iPAR molecules per aggregate whose mean value increases upon extracellular hyperosmotic stress.

Here we describe the design and construction of iPAR and the associated molecular cloning and bioimaging methodology and demonstrate the method’s utility to improvement the reliability of cytoplasmic protein aggregation investigations. We make iPAR openly accessible as a resource to the research community.

## 2. Material and Methods

### 2.1 Strains and plasmids used in the study

The yeast cell strains and plasmids used in this study are listed in Table 1 and 2, respectively and the oligonucleotides used in Table 3.

**Table 1:**
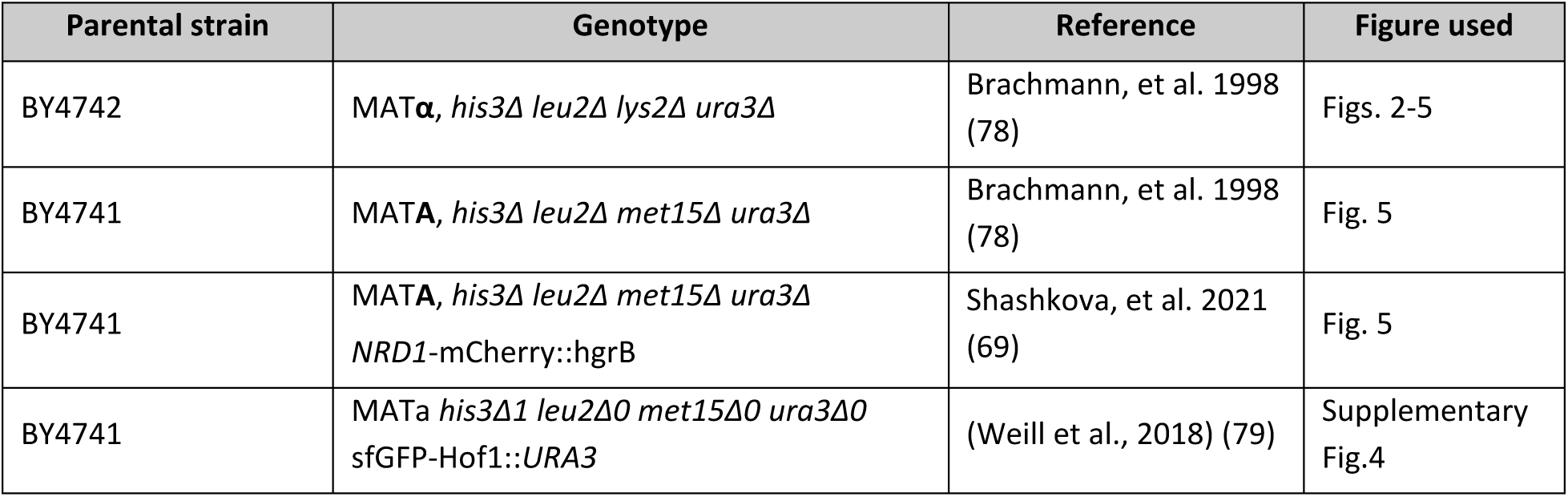
List of background yeast strains used in this study.

**Table 2:** List of plasmids used in this study.

| Plasmid | Genotype | Reference | Figure Used |
| --- | --- | --- | --- |
| pLS190 | pRS316 expressing ΔssCPY*-EGFP from the *PRC1* promoter | Stolz and Wolf, 2012 (44) | Fig. 2 |
| pCM695 | pRS316 expressing -GFP from the *CUP1* promoter | Laidlaw et al., 2021 (80) | Fig. 2 |
| pLS191 | pRS316 expressing ΔssCPY*-mEGFP from the *CUP1* promoter | This study | Figs. 2-5 |
| pLS195 | pRS316 expressing ΔssCPY*-mScarlet-I from the *CUP1* promoter | This study | Fig. 4 |
| pLS196 | pRS316 expressing ΔssCPY*-mNeonGreen from the *CUP1* promoter | This study | Fig. 4 |
| pCM264 | Mup1-EGFP from the *MUP1* promoter | MacDonald *et al.*, 2015 (81) | Supplementary Fig. 1 |
| pLS199 | pRS315 expressing ΔssCPY*-mEGFP from the *CUP1* promoter | This study | Not applicable |
| pLS200 | pRS315 expressing ΔssCPY*-mScarlet-I from the *CUP1* promoter | This study | Not applicable |
| pLS198 | pRS315 expressing ΔssCPY*-mNeonGreen from the *CUP1* promoter | This study | Not applicable |

**Table 3:**
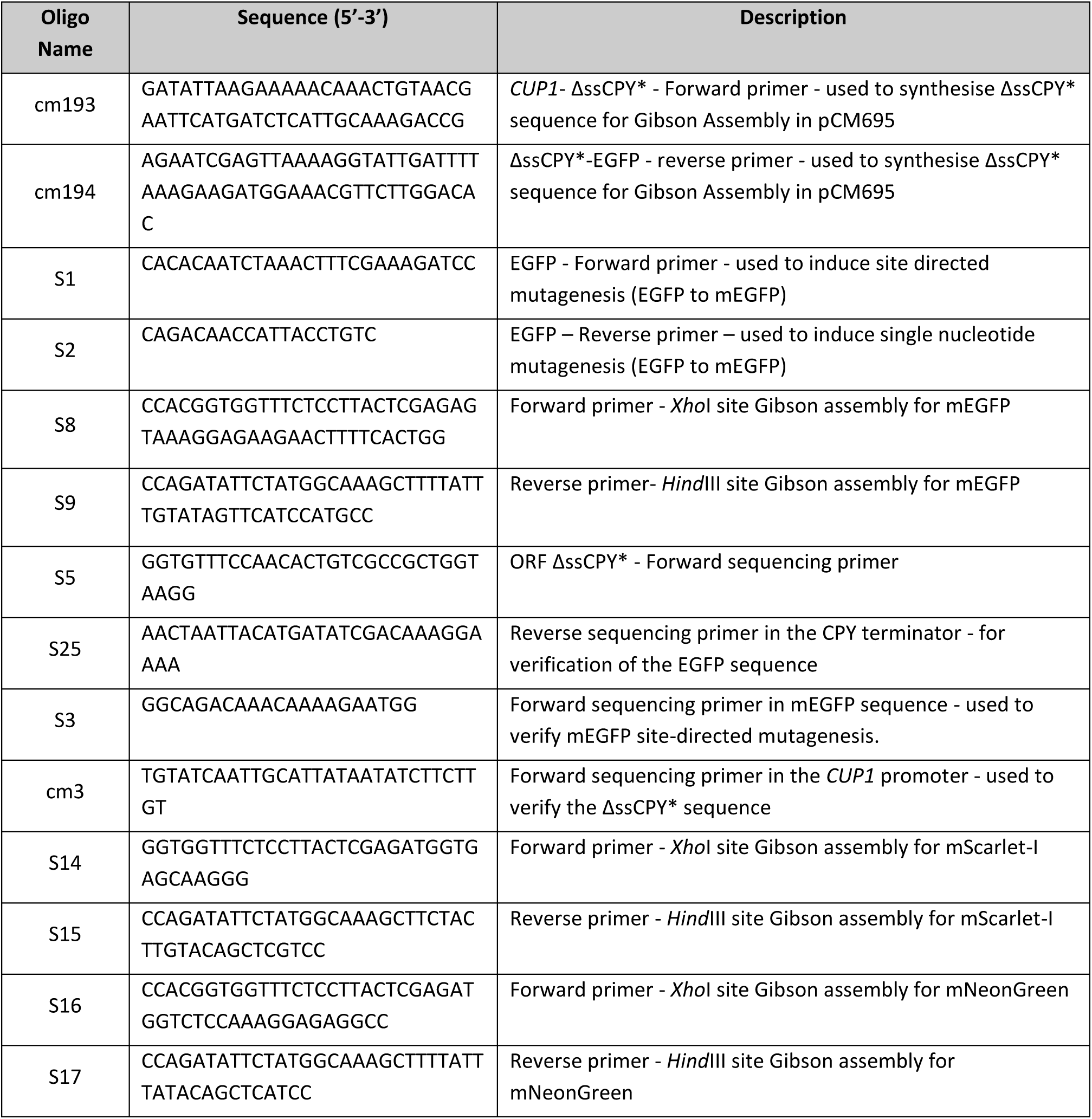
Primers used for the construction of the initial iPAR fusion construct *CUP1*-ΔssCPY*-mEGFP and subsequent variants using mScarlet-I and mNeonGreen fluorescent proteins.

### 2.2 Plasmid construction

An initial iPAR fusion construct *CUP1*-ΔssCPY*-mEGFP was generated using several cloning steps. Initially, the parent plasmid encoding *PRC1*-ΔssCPY*-EGFP (pLS190) was modified by site-directed mutagenesis using the S1 and S2 primers to incorporate the monomeric A206K mutation in EGFP (82). This template was then used to amplify ΔssCPY* (oligos cm193 and cm194) and mEGFP (oligos S8 and S9) with compatible regions for Gibson Assembly (83). ΔssCPY*-mEGFP was recombined between the *CUP1*-promoter and the *CYC1* terminator of pCM690 linearized with *Eco*RI and *Hind*III to generate pLS191 (C*UP1*-ΔssCPY*-mEGFP). The assembly strategy introduced 5’ *Xho*I and 3’ *Hind*III restriction sites flanking mEGFP. Plasmid pLS190 was linearized with *Xho*I and *Hind*III and Gibson assembly was used to exchange mEGFP with mScarlet-I (84) (using oligos S14 and S15, and using two PCRs separately to generate the *Xh*oI site) and mNeonGreen (85) (using oligos S16 and S17) variants of iPAR (respectively denoted as plasmids pLS195 and pLS196).

To maximise downstream applications of iPAR, in addition to creating red and green fluorescent variants with brighter fast maturing fluorescent proteins, we also switched the auxotrophic marker genes for plasmid selection (from *URA3* to *LEU2* selection). This was achieved by generating the *LEU2* gene from the integration plasmid pRS305, including ∼300bp of plasmid common to the ΔssCPY* reporter expression plasmid to facilitate recombination (based on pRS316). The PCR product was transformed into wild-type yeast alongside the *URA3* expression plasmid allowing the marker to be converted to *LEU2* by homologous recombination (see Supplementary Information).

### 2.3 Site-directed mutagenesis

The NEB Q5® Site-Directed Mutagenesis Kit (part number: E0554S, New England Biolabs Inc.) was used to perform the mutation responsible for mEGFP following the manufacturer’s protocol, with designed primers (S1 and S2, see Table 3) used at a concentration of 10 µM and the template DNA at a concentration between 1 to 25 ng/µl. The reaction mix was incubated for 5 min at room temperature before bacterial transformation.

### 2.4 Gel DNA extraction

To extract linearized plasmid backbones, gel DNA extraction was performed using the “QIAquick Gel extraction kit” (part number: 28706X4, QIAGEN, Ltd.), following the supplier’s instructions. In short, the DNA band of interest (cut from the agarose gel following electrophoresis) was transferred into a sterile 1.5 ml Eppendorf tube. QG buffer was added to the tube to dissolve the gel (at a 3:1 volume proportion) and incubated for 10 min at 50°C. The sample was loaded onto a silica-membrane-based spin column (1.5 ml volume) and centrifuged at 13,000 rpm. After discarding the supernatant, the column was rinsed once with 100% isopropanol followed by a wash with PB buffer. A final elution was performed by loading 50 µl of either EB buffer (10 mM Tris.Cl, pH 8.5) centrifuged at 13,000 rpm into a clean, sterile 1.5 ml Eppendorf tube.

### 3.5 Cell culturing

Single colony isolates from frozen stock following 24-48 h growth at 30°C were used to inoculate 5 ml liquid culture of either Yeast Extract-Peptone-Dextrose media (YPD: 2% glucose, 1% yeast extract, 2% bacto-peptone) or synthetic drop-out media lacking uracil (2% glucose, 1x yeast nitrogen base; 1x amino acid and base drop-out compositions (SD-URA, Formedium Ltd, UK), according to cell strains and selection requirements. Yeast cells were grown in the prepared liquid culture to mid-log phase (OD_600_ = 0.4-0.6) at 30°C before harvesting for imaging. A 100 mM copper sulphate stock solution was prepared, filter-sterilised with 0.22 µm diameter cut-off filters, and stored at room temperature. For the induction experiments, cells were first grown for 1-4 h in media containing 5 µM copper chelator bathocuproine sulfonate (BCS) before washing and incubation in media containing 100 µM copper sulphate to induce expression via the *CUP1* promotor (86). To promote the formation of aggregates, cells at the log phase were harvested, diluted to approximately OD_600=_ 0.2 and heat shocked for 2 h at either 37°C, 42°C or 30°C (the latter temperature being the control condition). The cells were then harvested and prepared for imaging with confocal microscopy.

### 3.6 Vacuole labelling

To label vacuoles, 0.8 µM FM4-64 (87) was added to 1 ml of cell culture in YPD-rich media and incubated with shaking for 1 h. Cells were then washed two times with SC media then grown for a further 1 h chase period in SC media lacking dye. After incubation, samples were prepared for imaging.

### 3.7 Sample preparation for imaging

Imaging was performed in “tunnel” slides (88) using 22×22 mm glass coverslips (No. 1.5 BK7 Menzel-Glazer glass coverslips, Germany). To immobilize cells to the surface, 20 µl of 1 mg/ml Concanavalin A (ConA) was added to the tunnel slide (89). Excess ConA was rinsed with 200 µl of imaging media before 20 µl of cells were added, incubated for 5 min upside down in a humidified chamber to promote cell adhesion. Finally, any unbound cells were removed by washing with 200 µl of imaging media and sealed with fast-drying nail varnish before loading on the microscope for imaging (90). Time-lapse experiments were performed in 35 mm glass-bottom dishes (Ibidi GmbH, Germany) with similar ConA coating methods adapted to the dishes support (91). 300 µl of 1 mg/ml of ConA were added to the dishes and incubated for 5 min then washed three times with sterile water. The dishes were then dried under a laminar flow hood ready for imaging. Typically, mid-log phase cells were diluted to OD_600_ <0.1 before addition to the ConA coated dish and incubated for 5 min at room temperature. The dish was washed two times with imaging media to remove any unbound cells and finally topped with fresh media for imaging.

### 3.8 Confocal microscopy imaging

Cell strains were excited using 488 nm and 561 nm wavelength lasers on the LSM 880 Zeiss microscopes with a 1.4 NA (Nikon) objective lens. Intensity and gain were optimised and then maintained for each experiment. Green fluorescence (from mEGFP and mNeonGreen fluorophores) was imaged using 2% laser excitation power and red fluorescence (from the mScarlet-I fluorophore) with 1% power to minimise photobleaching. Detector digital gain was set to 1 with a scanning time of 1.23 seconds per frame. Z stack images to generate 3D movies of cells expressing aggregates were acquired with 0.33 µm thick sections across the sample covering 5-6 µm thickness. FM4-64 vacuolar staining (87) was imaged with the 561 nm wavelength laser at 5% laser power using a bandpass emission filter range set to 578-731 nm. Timelapse imaging was performed by acquiring 10 min intervals of 3 μm thick section slices images over 90 min for optimal cytoplasmic volume visualisation during cell division (as described in previous work (92)).

### 3.9 ImageJ image analysis

Confocal microscopy data were analysed using ImageJ/Fiji software (ImageJ 2.14.0/1.54f/Java 1.8.0_322) to extract fluorescence intensities from pre-defined segmentation outlines. Cell outlines were generated either manually using the ImageJ selection tool or in a semi-automated process using the `Cell Magic Wand` plugin (93). Fluorescent foci within each cell were detected using our bespoke ImageJ macro `SegSpot` allowing for the selection of a threshold method (within the range of inbuilt thresholding functions available in ImageJ) and object detection function within pre-defined cells outlines or regions of interest stored in ImageJ `ROI Manager`. Finally, pixel intensities and area parameters of the identified foci were extracted and displayed in an output table (See Supplementary Figure 5). Z stack images were visualized with the `3D project` inbuilt ImageJ plugin.

### 3.10 Slimfield microscopy

Preliminary attempts to measure the mobility of iPAR-labelled aggregates using fluorescence recovery after photobleaching (FRAP) were technically challenging due likely to their relatively high diffusion rates. Because of its single-molecule precise detection sensitivity and rapid millisecond sampling capability, we used Slimfield microscopy (61, 66, 67, 77, 94–96) to characterise the iPAR-labelled aggregates in terms of their molecular stoichiometry (defined as number of fluorescent iPAR tags estimated per distinct fluorescent focus detected) and their mobility within the cell cytoplasm (in terms of the effective diffusion coefficient of tracked iPAR foci). This method enables quantification of the spatial dependence of rapid diffusion *in vivo* in ways that more traditional technologies such as fluorescence correlation spectroscopy (FCS) cannot. Cells expressing iPAR were imaged using excitation via an epifluorescence narrowfield laser beam (96) to generate a Slimfield profile with wavelength 488 nm (Obis LS laser) set to 20 mW power at the sample using 1,000-1,500 frames per acquisition at 5 ms per frame sampling time.

Aggregates were produced following the established standard condition, and cells grown to log phase were induced for iPAR expressing using 100 µM copper for 2 h including 1 h heat shock at 37 °C. Osmotic stress with 1 M NaCl and 1.5 M sorbitol was applied and compared to the control condition with cells in 50 mM NaPi.

Protein aggregates were tracked using our in-house software platform which could be implemented in both MATLAB (97) and Python (98) modalities, which uses iterative Gaussian fitting (99) to pinpoint the spatial location of tracked fluorescent foci in complex live microbial cells to approximately 40 nm lateral precision and quantifying stoichiometry, copy number and mobility parameters (100). Stoichiometry was determined by normalising the initial unbleached track intensity with the brightness value estimated for a single iPAR molecule *in situ* in live cells and from *in vitro* experiments on purified iPAR molecules using step-wise photobleaching, then rendering stoichiometry distributions using kernel density estimation analysis (101).

Diffusion coefficients were estimated from the initial gradient of the mean square displacement versus time interval relationship generated for each track (102, 103), assuming the solution environment is purely viscous as opposed to viscoelastic (104).

## 3. Results

### 3.1 Construction of an inducible monomeric marker for cytoplasmic aggregation in budding yeast cells

The vacuolar hydrolase CPY traffics through the biosynthetic pathway as an inactive precursor before activation in the yeast vacuole (105). A mutant version of CPY prone to aggregation, denoted CPY* (44), has been used in previous studies as a model to assess protein folding and regulatory control of misfolded proteins (105–108). The CPY* variant carries a single amino acid substitution of glycine for arginine at position 255 (G255R) near the enzymatic active site (Figure 1.A – 1.C). Deletion of the N-terminal signal peptide (Δss) of CPY inhibits entry to the secretory pathway and consequently the hydrolase mislocalises to the cytoplasm (109). The ΔssCPY* mutant, which aggregates in the cytoplasm, serves as a useful marker for protein aggregation (50, 51, 110, 111). However, the endogenous *PRC1* promotor (112) typically used to induce expression of this aggregate marker is metabolically regulated (54, 55); therefore, expression, and aggregation, often vary depending on the specific growth and stress conditions resulting in potential difficulties of interpretation.

To overcome this limitation, we generated a fusion construct which expressed ΔssCPY* from the copper inducible *CUP1* promoter (113) in the presence of 100 µM copper sulphate (see Methods and schematic Figure 2.A), using definitive monomeric fluorescent protein tags (monomeric EGFP in the first instance) to mitigate against issues associated with fluorescent protein oligomerization. Using a titration from 0 - 200 µM copper sulphate on a GFP-tagged methionine permease we previously used for membrane trafficking studies (81). We have routinely used the *CUP1* promoter because under basal media conditions, which have a very small amount of copper, expression levels are low. Although expression can be further reduced with copper chelation, this also inhibits cellular growth(114). Therefore, we use media lacking copper to culture cells to appropriate log phase and density for experiments, before adding up to 100 µM copper to robustly induce expression. However, 100 µM copper has no detectable phenotype on cellular process we have measured. Furthermore, we confirmed that copper had no measurable effect on fluorescence levels. Flow cytometry was used to define background fluorescence in wild-type cells and distinguish fluorescence of Mup1-EGFP expressing cells (Supplementary Figures 1 and 2).

**Figure 2:**
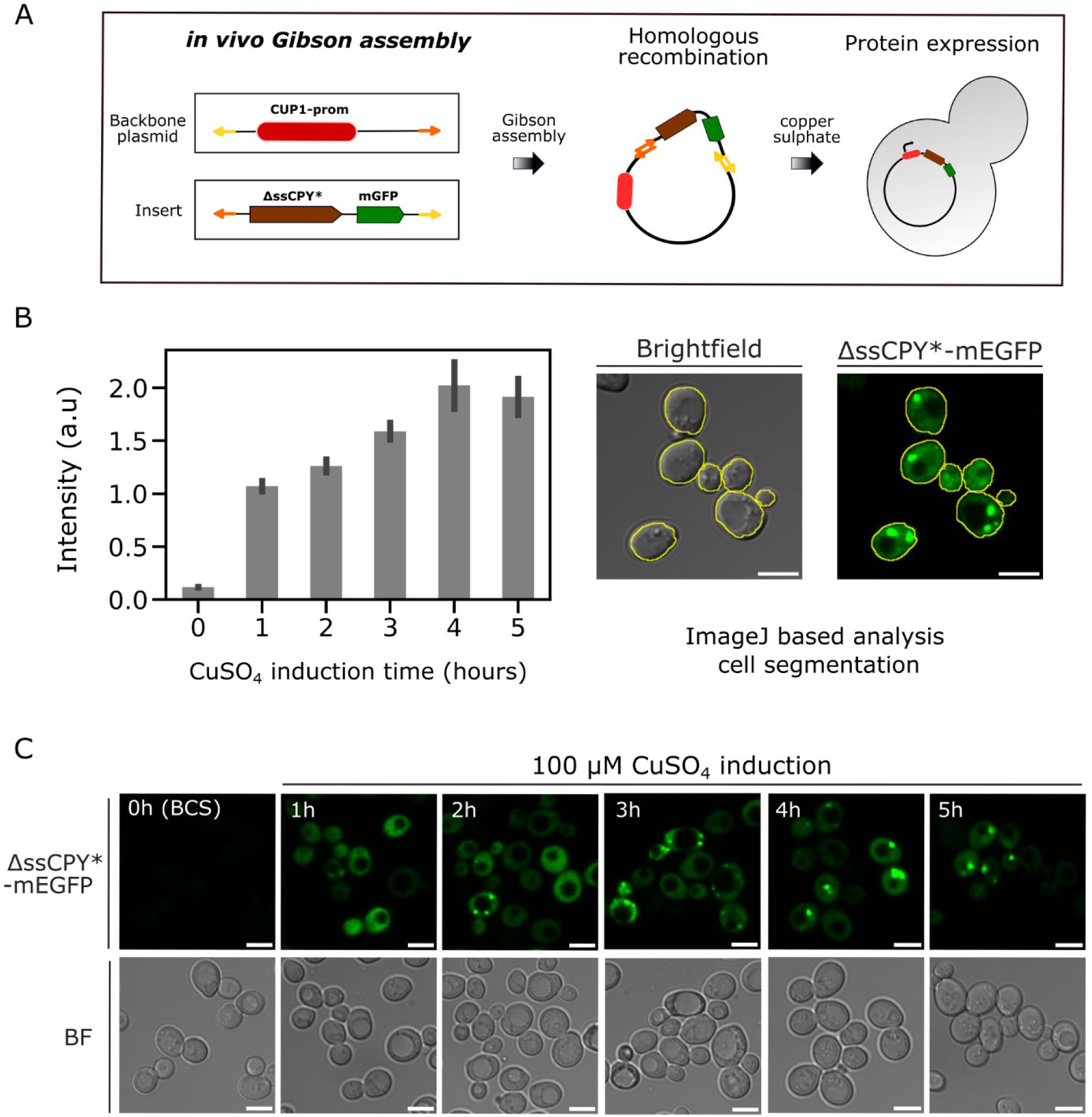
Induction of *CUP1* promoter by copper sulphate results in expression of protein aggregates, visible in confocal microscopy. A) Schematic representation of cloning strategy to produce copper-inducible cytoplasmic ΔssCPY*-mEGFP aggregates. B) Bar plot for the fluorescence intensity of *CUP1*-ΔssCPY*-mEGFP incubated in the copper chelator BSC (0 h) or following induction by 100 µM copper sulphate, at 1 h, 2 h, 4 h and 5 h, n = 100 cells for each condition, s.e.m. error bars represented. The micrographs on the right show cell segmentation using the `Cell Magic Wand` ImageJ tool applied to brightfield images. These segmented images were then used to quantify the total fluorescence intensity from the GFP channel corresponding to each cell. C) Fluorescence micrographs representing the ΔssCPY*-mEGFP aggregation at different induction time points.

Copper-dependent expression levels of *CUP1*-ΔssCPY*-mEGFP in budding yeast cells were characterised using confocal microscopy. Induction times from 1 - 5 h were used followed by imaging and subsequent image segmentation analysis to extract the fluorescence intensity and the integrated pixel volume information of cells and protein aggregates. We found that expression of iPAR could be rapidly induced in the presence of 100 µM copper sulphate (Figure 2.B), with a strong increase observed after 1 h copper exposure, with an apparent slowing after 2 h and 3 h exposure and steady-state expression levels after approximately 4 h (Figure 2.B and C). At 5 h induction we noticed a small decrease in fluorescence intensities, which was consistent with the activity of clearance pathways associated with protein aggregation.

A 2 h copper incubation time was selected as a standard induction condition to express the ΔssCPY*-mEGFP marker to generate a sufficient pool of protein aggregates for subsequent analysis. We noticed that after 2 h expression there was a reasonable level of expression and several aggregates forming in the cytoplasm (Figure 2.C).

We then characterised the effect of temperature on cells expressing ΔssCPY*-mEGFP following heat shock. As expected, cells grown for 1 h at 30°C exhibited very few protein aggregates, however, shifts to heat stress conditions using temperatures of 37°C or 42°C resulted in measurable iPAR aggregate formation (Figure 3.A). There was a significant increase in cells following heat shock at both 37°C or 42°C in comparison to any cells at 30°C that had detectable aggregates of ΔssCPY*-mEGFP (Figure 3.B). A significant increase in number of aggregates was observed, in addition to the number of cells in which aggregates were detected, following heat stress (Figure 3.C).

**Figure 3:**
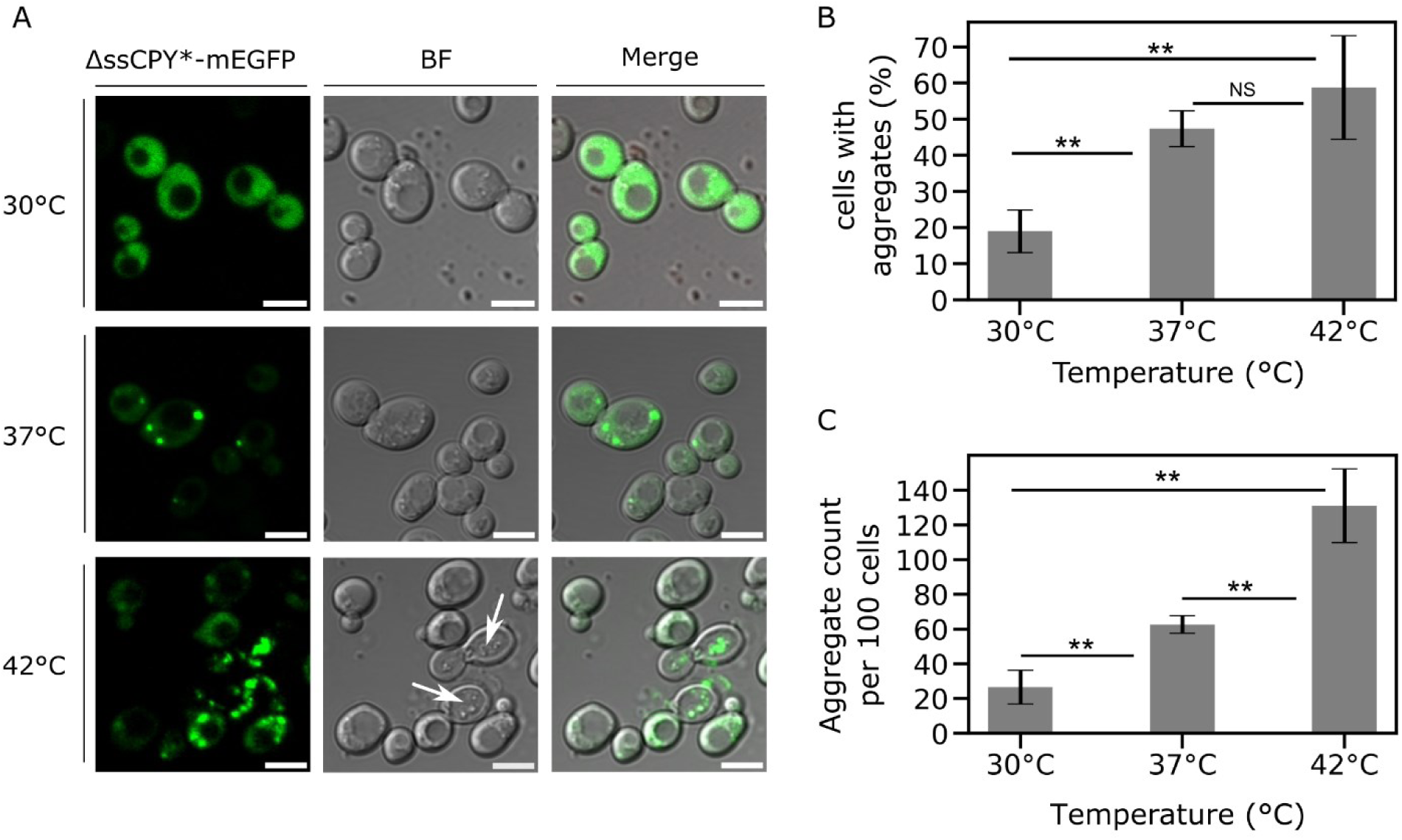
Short-term heat shock induces the formation of aggregates. A) Confocal micrographs from a representative cell population of yeast cells expressing the *CUP1*-ΔssCPY*-mEGFP protein after induction with copper sulphate for 2 h followed by 1 h at either the initial growth temperature 30°C or the heat shock temperatures of 37°C and 42°C. White arrows indicate dead cells in the brightfield channel, which were not used in subsequent analysis. Scale bar: 5 µm. B) Bar plot representing the percentage of cells which were positive for aggregates for cells exposed to the control 30°C, or the 37°C and 42°C heat shock. Non-significance is indicated by a Student’s *t*-test p value ≥0.05, the double asterisk indicates a p value <0.05. C) Bar plot showing the number of aggregates detected and counted in the cell population, bringing it to n = 100 cells in total, s.d. = error bars. See also Supplementary Table 2.

Between 30°C and 37°C, we observed an increased number of aggregate-positive cells (defined as a cell which contains at least one detected iPAR fluorescent focus) by a factor of approximately 2.5, from an average of 19% (±5.8, s.d.) to 47% (±9.4), corresponding to a Student’s *t*-test p value of 7.59 × 10^−5^ (i.e., highly significant). Similarly, between 30°C and 42°C, the pool of aggregate-positive cells increased by a factor of approximately 3 from 19% (±5.8) at 30°C to 59% (±14.3) at 42°C with a significant p value of 5.00 × 10^−3^. Although 42°C induced a greater number of aggregate foci across the population, we also detected elevated levels of cell death (Figure 3.A; arrows). Additionally, there was no significant increase in aggregate-positive cells by heat shocking at 42°C compared with 37°C (p = 0.261) (see Figure 3.B and Supplementary Table 1).

The total number of detected aggregates increased by a factor of 2.4 from 30°C to 37°C, and by a factor of 4.9 between 30°C and 42°C; and, although the number of aggregate-positive cells was similar between 37°C and 42°C, we still observed a significant increase in the number of aggregates detected (Figures 3.B,C and Supplementary Table 2). We subsequently used 2 h copper induction followed by 1 h heat shock at 37°C as our standard protocol, which we found to be sufficient to induce trackable ΔssCPY*-mEGFP aggregates without compromising the phenotype or viability of the cells.

To expand the utility of the iPAR reagent, the mEGFP fluorescent tag was flanked with unique cutting sites (5’ *Hind*III and 3’ *Xho*I sites) to enable interchangeability and future extension of the construct library for DNA insertion to encode different fluorescent proteins (Figure 4.A). We used this strategy to create iPAR variant *CUP1*-ΔssCPY*-mNeonGreen and *CUP1*-ΔssCPY*-mScarlet-I, which we found also formed inducible aggregates following the optimised protocol described above in a qualitatively similar manner (Figure 4.B).

**Figure 4:**
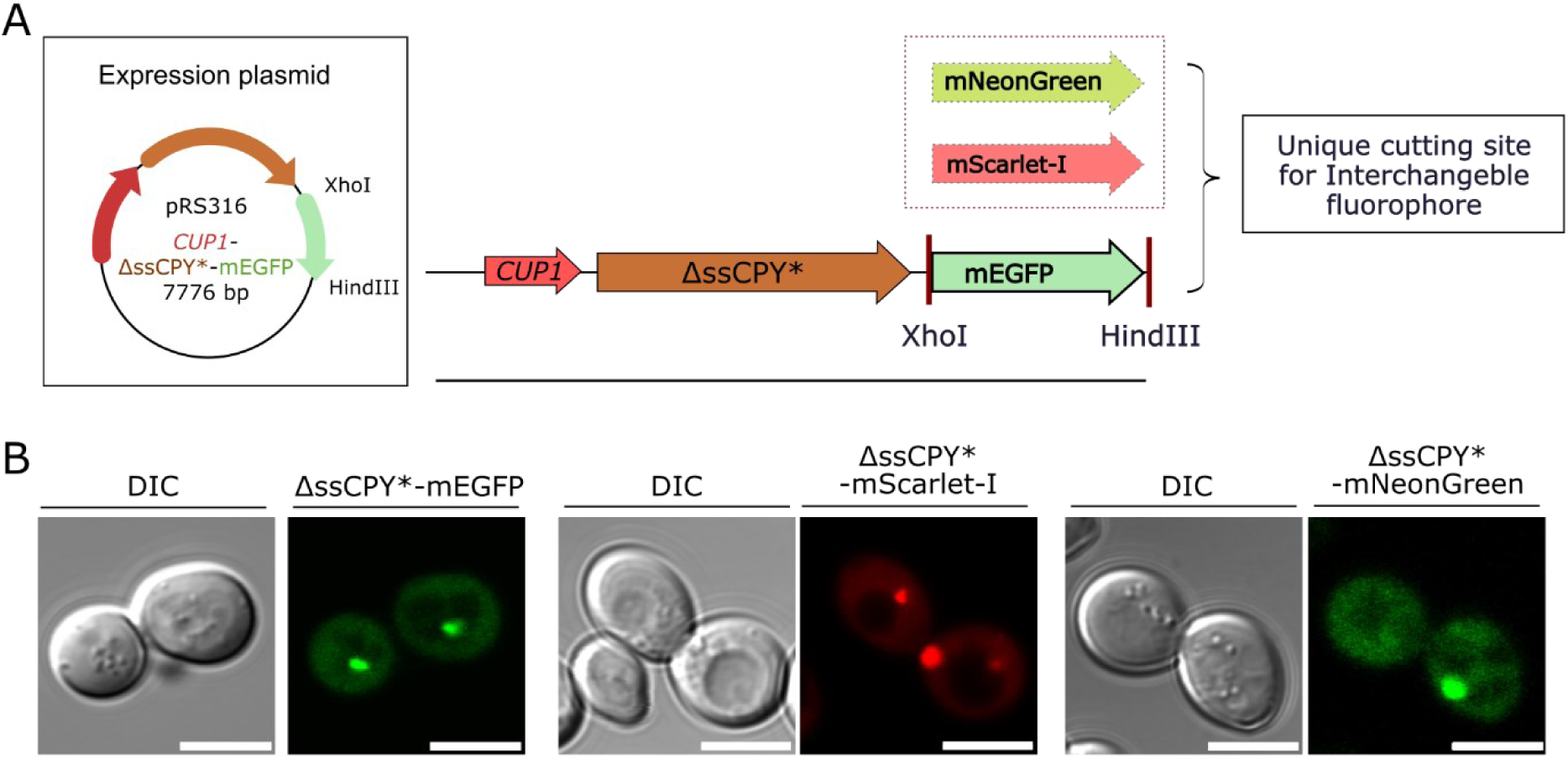
iPAR enables interchangeable monomeric fluorescent proteins to be used for reporting on protein aggregation inside the cytoplasm of living yeast cells. A) Schematic of the expression plasmid constructed for *CUP1*-ΔssCPY*-mEGFP, the fluorophore with *Hind*III and *Xho*I cutting sites used to facilitate the exchange of fluorescent markers. B) From left to right, micrographs with differential interference contrast (DIC) and fluorescence channel for *CUP1*-ΔssCPY* in pRS316 with the mEGFP, mScarlet-I and mNeonGreen fluorescent proteins shown respectively.

### 3.2 Cytoplasmic aggregates and localisation in time and space in budding yeast

We performed further characterisation of iPAR to focus on spatiotemporal dynamics of newly formed aggregates. We first investigated the number of aggregates and their spatial distributions between mother and daughter cells. Figure 5.A shows the analysis focused on budding cells, where mother and daughter cell images were independently segmented using our bespoke `SegSpot` macro coded for ImageJ which enabled thresholding and object detection of fluorescent foci (see Methods and Supplementary Figure 4). The area and intensity of fluorescent foci were automatically extracted by this macro, and their values plotted (Figure 5.B). Jitter plots revealed that the mean foci areas measured in mother cells were approximatively twice as large as those measured in daughter cells, with a mean focus area of 0.99 (±0.74) µm^2^ measured in mother cells v*s* 0.39 (±0.29) µm^2^ for daughters (Figure 5.B: left plot and Supplementary Table 3). Mother cells contained aggregates of higher volume with a mean fluorescence intensity significantly higher than daughter cells, corresponding to an integrated intensity (measured in arbitrary units A.U., rounded to nearest 100 A.U.) of 52,400 A.U. (±8,500) *vs* 37, 100 A.U. (±9,800) respectively (right plot of Figure 5.B, see also Supplementary Table 3). We note that the distribution of numbers of aggregate foci in both cell types is heterogeneous but more pronounced in mother cells (Figure 5.B), which was also reflected by higher standard deviation values. These results suggest a polarity behaviour of formation/clearance of ΔssCPY*-mEGFP during cellular growth resulting in statistically different sizes of aggregates between two cells which are dividing (the older cells displaying larger aggregates with higher intensities than those of the emerging daughter buds).

**Figure 5:**
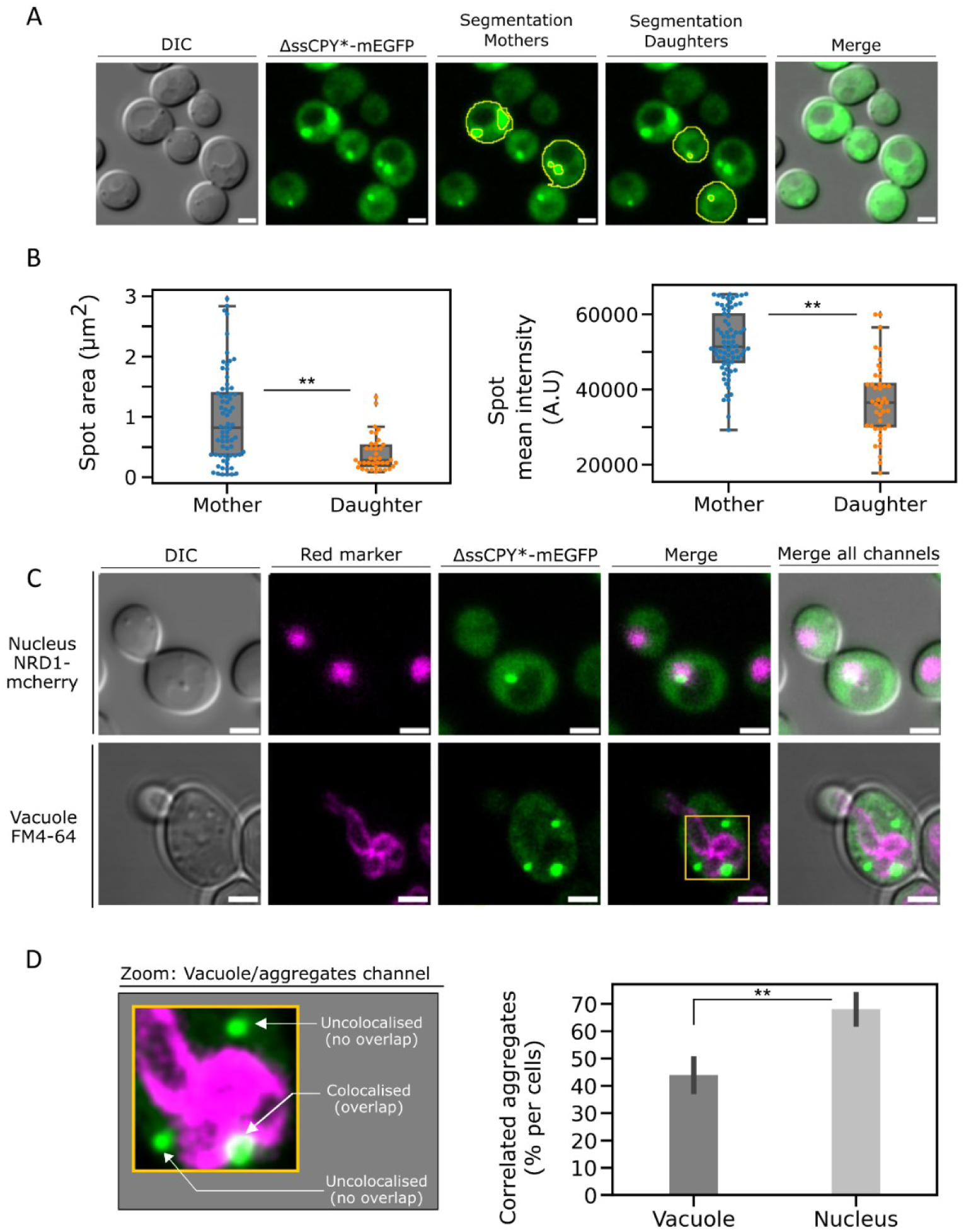
Protein aggregates localise specifically to vacuolar and nuclear compartments. A) Semi-automated segmentation (a combination of the ImageJ selection tool and our bespoke automated macro processing) of mother cells and daughter cells to characterize fluorescent foci. From left to right: DIC image of the cell, fluorescence channel, segmentation of the mother cells, of the daughter cells and merge of the fluorescence channel with the DIC. Scale bar: 2 µm. B) Characterization of aggregate foci, jitter plot of the detected foci area between mother cell and daughter cells. On the right, jitter plot of the intensity measured in each fluorescent focus identified. Outlier detection and removal was performed using standard interquartile methods (115, 116). C) Fluorescence micrographs of dual label strain for simultaneous observation of aggregates and key cellular compartments. Top row shows the nucleus labelled by nuclear reporter Nrd1-mCherry background strain, bottom row shows the vacuole labelled with FM4-64 (87), which mark the vacuole location. Micrographs showing the brightfield, the red channel with the marked compartment of interest, the green channel with the iPAR aggregate reporter and the merge of both fluorescence channels along the brightfield. Scale bar: 5 µm. D) (left) Zoom-in of region highlighted in panel C with (right) estimate of the percentage of detected aggregates which are either colocalised with the vacuole or nucleus compartments (s.d. errorbars, n=100 cells, the double asterisk indicates a p value <0.05).

We then sought to verify whether iPAR indicated any qualitatively similar spatiotemporal behaviour as reported previously for other cytoplasmic aggregation reporters (53, 117, 118). For example, ΔssCPY* aggregates were previously shown to be localized in JUNQ and IPOD (119) inclusion bodies, observed near the nucleus (120) and the vacuole (121), respectively. To elucidate whether our induced ΔssCPY*-mEGFP colocalised near the membrane of either of the nucleus or the vacuole, we constructed dual colour cell strains including a fluorescent red tag as a reporter for the location of the nucleus or the vacuole. Figure 5.C shows the resulting dual colour images of representative live cells, the top row showing Nrd1-mCherry (59, 122) marking the nucleus, the bottom row showing using FM4-64 pulse-chased labelling to mark the vacuole (see Methods), both simultaneously expressed with ΔssCPY*-mEGFP.

We quantified the proportion of aggregates present in each cellular compartment, by assessing the proximity/colocalization of both colours (micrographs in Figure 6.D) and found that a mean of approximately 44% of aggregates colocalised with the vacuole compartment and 68% with the nucleus (Figure 5.D). This result is broadly consistent with earlier observations that a significant number of aggregates appear to localise both near the nucleus or vacuole (93). The higher percentage of aggregates identified as being associated with the nucleus may indicate that aggregates preferentially sequestrate into JUNQ inclusion bodies.

**Figure 6:**
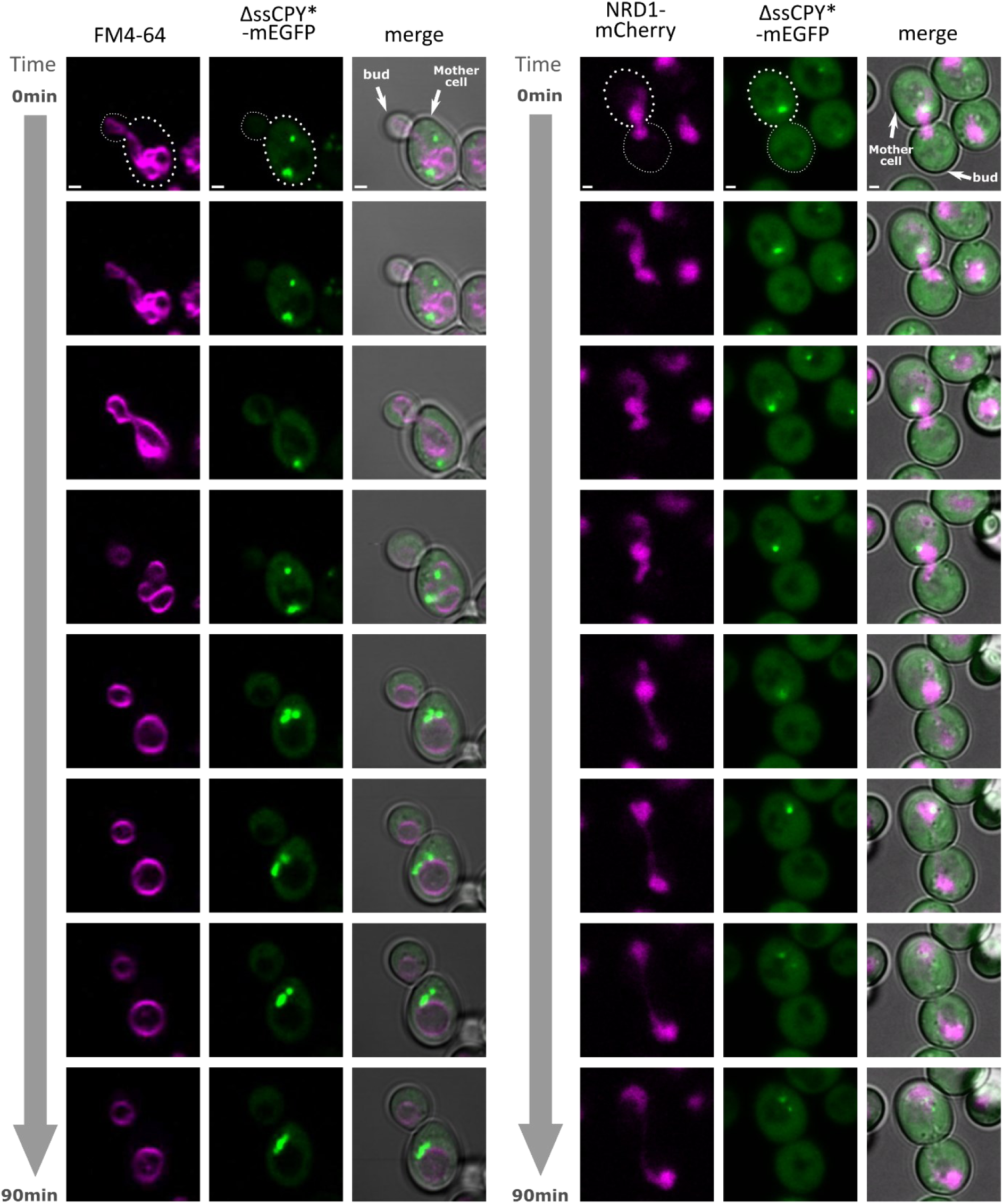
Protein aggregates are localized near to the vacuole and nucleus during cell division. Cells expressing the ΔssCPY*-mEGFP trackable aggregates (generated after 2 h copper sulphate induction including 1 h heat shock at 37°C) in combination with either Nrd1-mCherry expressed in the nucleus or a WT background strain labelled with FM4-64 (87) at the vacuole, imaged using confocal microscopy over 90 min during cell division. Micrographs show the red channel for those two markers of interest, the green channel of the imaged aggregate marker and the merge of both fluorescence channels along the brightfield. White arrows indicate the mother cell and the bud position. Scale bar: 1 µm.

We also acquired 3D data to visualise the patterns of aggregate spatial expressions inside the entire volume of the cell (Supplementary Figure 3 and Supplementary Videos 3-6). 3D projections of cells expressing iPAR, including labels of either the vacuole or nucleus, further confirmed the presence of cytoplasmic aggregates, appearing preferentially in the mother cells and confirming localisation in regions that are in likely contact with the nucleus and vacuole membrane to within our optical resolution limit of approximately 250 nm.

Finally, we performed time-course experiments during cell division with the dual label strains detailed above. In both cases, as a cell divides, we observed protein aggregates sequestrated in the mother cell (Figure 6 and Supplementary Videos 1 and 2). We observed that both vacuoles and nuclei were inherited into budding daughter cells whilst aggregates were retained in the mother cells. We note both events occurs at different stage of the cell cycle, the vacuole is inherited at early stages (∼20 min) of the budding process but the nucleus is one of the last (∼60 min) (123).

This observation reinforces the hypothesis that there is a diffusion barrier between mother and daughter cells maintained during cell division (123–125). The sequestration of misfolded cytoplasmic proteins has been reported previously as being a highly conserved quality control process which is crucial to cellular rejuvenation (126–130); the presence of ΔssCPY* associated with both JUNQ and IPOD inclusion bodies suggests a potential cellular recognition and cellular response for clearance and degradation.

### 3.3 Using iPAR in conjunction with Slimfield to quantify the molecular stoichiometry of aggregates and their spatial distribution and mobility in live cells

We used Slimfield on live cells expressing the mEGFP iPAR variant to enable us to the count how many iPAR molecules are present in aggregates and how rapidly aggregates diffuse inside cells (Figure 7.A). Cells were visualised in normal 50 mM NaPi imaging buffer, as well as 50 mM NaPi supplemented with either 1.5 M NaCl or 1 M sorbitol, typical conditions to induce hyperosmotic stress; NaCl and sorbitol are both crowding agents of distinct nature therefore with a different potential of interaction on metabolic functions and oligomerisation(131).

**Figure 7:**
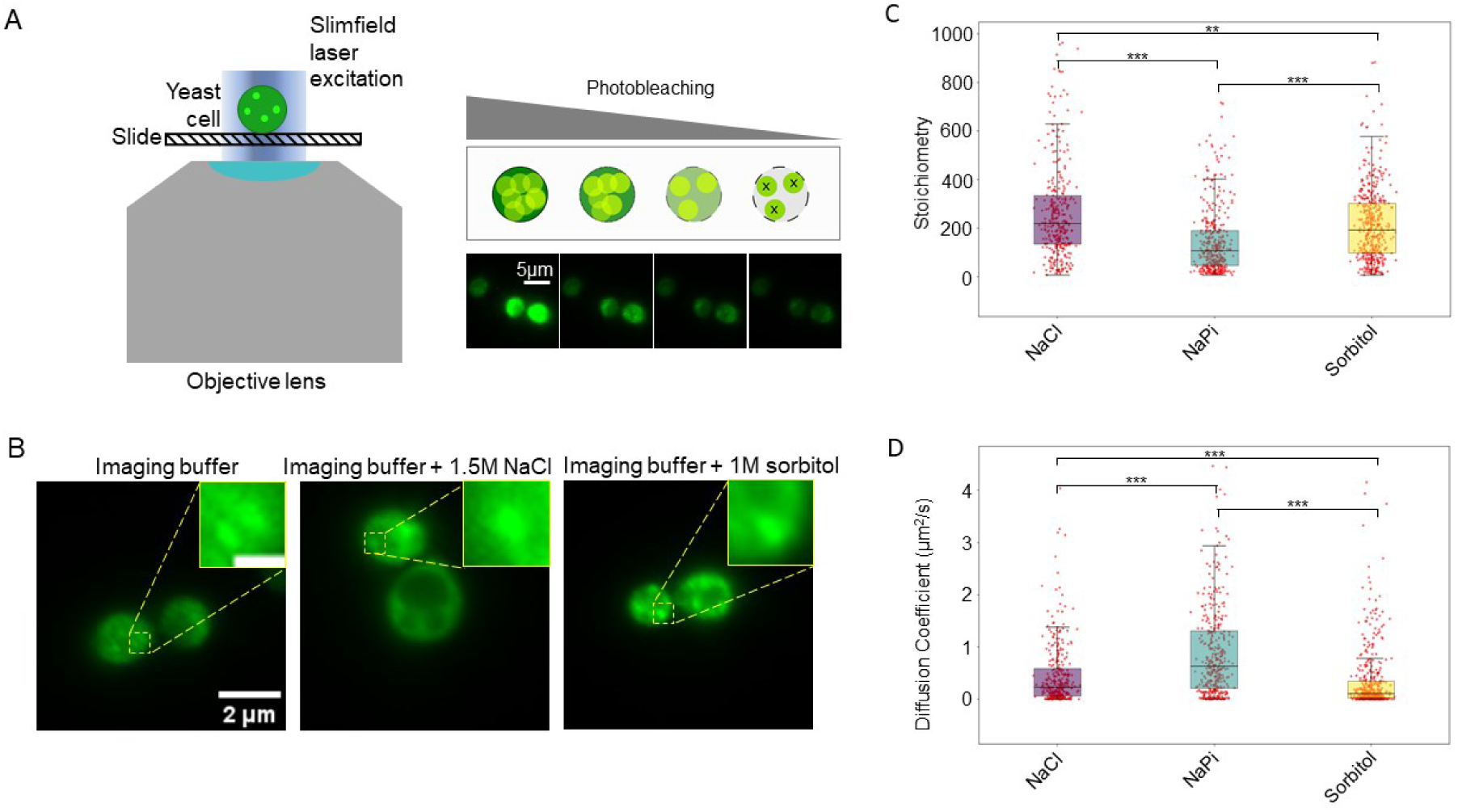
iPAR labelling is compatible with single-molecule precise millisecond timescale Slimfield microscopy. A) (left) cartoon representation of Slimfield excitation, in which the width of the laser beam is only a little larger than the diameter of a single cell, utilising the associated increased laser excitation intensity to enable detection of single iPAR molecules above the camera detector noise; (right) schematic representation of photobleaching of iPAR molecules inside cells which enables the detection of single molecules due to the subsequent increased mean spatial separation of remaining unbleached iPAR molecules, also visualised using yeast expressing ΔssCPY*-mEGFP. B) Representative images of yeast expressing ΔssCPY*-mGFP in normal imaging buffer, or under hyperosmotic stress in the form of 1.5 M NaCl or 1 M sorbitol respectively, insets showing distinct aggregate foci (inset scale bar 500 nm). C) Comparison of aggregate stoichiometries, and D) diffusion coefficients, under the previously mentioned stress conditions using box plots indicating the median value and interquartile range, with the aggregate populations in each condition showing statistically significant differences when compared using a Mann Whitney U test (for stoichiometries the corresponding p values (1 d.p.) are: NaCl:NaPi=1.1 × 10^−23^, NaCl:sorbitol=1.1 × 10^−3^, NaPi:sorbitol=1.4 × 10^−14^; for diffusion coefficients the corresponding p values are: NaCl:NaPi=3.3 × 10^−14^, NaCl:sorbitol=7.0 × 10^−8^, NaPi:sorbitol=7.3 × 10^−33^). Number of tracked foci for NaCl n=337, NaPi n=393, sorbitol n=430.

Slimfield images exhibited distinct fluorescent foci corresponding to protein aggregates (Figure 7.B), qualitatively similar in appearance to those observed with confocal and epifluorescence microscopy, which could be pinpointed using our bespoke localisation microscopy tracking software, optimised in budding yeast cells to a lateral spatial precision of approximately 40 nm (132). This analysis software enabled measurement of molecular stoichiometry of each tracked aggregate by using a method which converts their quantified integrated pixel brightness into the number of photoactive iPAR molecules utilising a stepwise photobleaching protocol (19) to determine the brightness of a single fluorescent protein molecule (88).

We observed an increase in stoichiometry for both of the hyperosmotic stress conditions applied, from a mean of 157 (±25) molecules per aggregate for the non-stress condition with cells in 50 mM NaPi buffer to 290 (±28) for 1 M NaCl (osmolarity equal to 2 osmol/L) corresponding to an 85% increase, while the stoichiometry measured for 1.5 M sorbitol (osmolarity equal to 1.5 osmol/L) was 217 (±17), a 38% increase compared to the control condition (Figure 7.C). The tracking software also enabled estimates of the lateral diffusion coefficient for each aggregate, indicating an associated reduction of aggregate mobilities in a hyperosmotic extracellular environment, consistent with an associated increase in intracellular molecular crowding (92). The control condition shows a diffusion coefficient of 0.99 (±0.15) µm^2^/s compared to 0.47 (±0.04) for 1 M NaCl and 0.36 (±0.06) µm^2^/s for 1.5 M sorbitol, corresponding to a decrease of 48% and 36% respectively. This quantitative analysis exemplifies iPAR being used in conjunction with an example of rapid single-molecule bioimaging tecnhology, Slimfield. It robustly quantifies differences of aggregation due to different hyperosmotic stress factors, for example the effect on aggregate stoichiometry and diffusion is of a greater extent when induced by 1 M NaCl salt exposure than for 1 M sorbitol, consistent with simple colligative differences is osmolarity.

It reveals a broad range for both stoichiometry and diffusion coefficient for aggregates, an observation which resonates with the concept of aggregate formation being driven by dynamic and heterogeneous protein nucleation inside cells. These observations indicate that these extracellular hyperosmotic environments bias the likelihood of protein nucleation events that result in aggregate formation.

More generally, these findings show that iPAR is compatible with high-precision rapid single-molecule localization microscopy using different osmotic stress factors to study protein aggregation in live cells.

## 4. Discussion

We have developed iPAR, an improved reporter for high-precision quantification of cytoplasmic protein aggregation in the budding yeast *Saccharomyces cerevisiae*. By replacing the metabolically regulated *PRC1* promoter with the copper sulphate inducible *CUP1* promoter and introducing definitively monomeric fluorescent tags, iPAR enables precise control of protein expression in growing cells with reduced interference from the fluorescent tag in the aggregation process. These modifications offer an alternative choice of reporter for stress-related studies and for investigating the dynamics of protein aggregation, compared to heat shock protein biomarkers of aggregation which use non-monomeric GFP (115). As with all fluorescent protein probes the mEGFP used in iPAR will have a maturation time. In the context of the experiments described here ΔssCPY*-mEGFP is expressed for 2 hours before imaging which provides ample time for maturation of mEGFP (∼22 mins (59, 88), however this maturation time could prove limiting in experiments that require the immediate imaging of iPAR upon expression. It should also be noted that any stoichiometry measurements of ΔssCPY*-mEGFP within aggregates are likely to be an under representation (in the region of 7% (59)) of the true number due to the presence of dark constructs that are non-photoactive. AS proof-of-concept, we used 1M NaCl and 1.5M sorbitol to induce different levels of cellular hyperosmolarity, though interesting future work could titrate these respective levels to compare phenotypic responses of these different osmolytes but at comparable osmolarities.

We first characterised iPAR by measuring the expression response of ΔssCPY*-mEGFP to 100 µM copper sulphate, indicating that a 2 h standard induction time was optimal to produce a strong fluorescence signal of protein aggregates. We then tested the effects of heat shock on aggregation following inducible expression. At 37°C, we measured a strong increase in aggregate-positive cells (greater than twice as many cells that contain protein aggregates compared to cells incubated at the 30°C no-stress control condition). At 42°C, we observed a similar number of aggregate-positive cells, but we detected a higher total number of aggregates across a population of cells as well as a higher number of aggregates per cell. However, the physiological cell phenotype of 42°C was visibly impaired in several instances, including abnormal morphology and dead cells, consistent with cell metabolic malfunction resulting in an increase in cytoplasmic aggregation. Therefore, we did not select this temperature in subsequent investigations using iPAR. A concentration of 100 µM copper sulphate was sufficient to induce aggregate formation and not to generate cellular defects from copper toxicity; future work in titrating different concentration levels of copper sulphate and observing function responses regarding aggregate properties could be valuable.

We verified that induced aggregates localise to the nucleus and vacuole JUNQ and IPOD compartments respectively, as reported from previous studies using existing aggregation reporters. We performed time lapse confocal microscopy imaging to quantify the extent of inheritance of the vacuoles and nuclei during asymmetric cell division of iPAR yeast cells in real time, showing directly on a cell-by-cell basis that these intracellular organelles are inherited to daughter cells whilst proteotoxic aggregates are retained in the mother cell (see Figure 6 and Supplementary Videos 1 and 2). These time-resolved observations taken using the same individual cells are consistent with earlier reports using separate imaging of organelles and aggregates across several different cells (118, 126), however, this is to our knowledge the first direct observation that such aggregates which appear to be associated with specific organelles are, unlike the organelles themselves, not inherited.

In budding yeast cells, the presence of multiple inclusion bodies typically observed during osmotic stress were shown previously to be further sequestrated in targeted cellular locations (118, 133). Aggregates may be actively recognized by cells and sequestrated in the mother cell volume, additionally, physicochemical properties such as local viscosity (134) and the molecular crowding at the junction between the two cells can potentially influence aggregate localisation, as suggested by the results of our previous study (92) on the investigation sub-cellular crowding dynamics. This molecular crowding at the junction between two cells may hold a key as to why these toxic aggregates are not inherited alongside their associated organelles. Experiments utilising iPAR with high-precision Slimfield measurements probing this putative junction effect may be valuable future experiments to address this hypothesis since, as we demonstrate here, Slimfield has the capability to robustly quantify the spatiotemporal dynamics of iPAR aggregates, showing that they are mobile inside cells and are comprised from as few as a few tens of molecules up to several hundred, whose mean value increases with extracellular hyperosmotic stress.

Slimfied microscopy enables rapid tracking of aggregates over the entire cell, provided they are within the ca. 1 micron depth-of-field of the microscope and have with sufficient contrast against background noise for detection. This enables measurement of the spatial dependence of rapid molecular mobility. Other measurement approaches for quantifying molecular mobility could in principle be used, for example FRAP and FCS, however the relatively slow scanning speeds currently prohibit easily reproducible measurements of molecular mobilities in different regions of the same cell at the same point in time. There are also a suite of different fluorescent-based super-resolved single-particle tracking approaches which could be used in complement to Slimfield (135, 136), though unless specific efforts are made to adapt these the sampling times which are possible are slower than Slimfield’s rapid sub-ms capabilities. It should also be noted, that Slimfield uses a localization microscopy approach which can pinpoint single molecular assemblies that span effective diameters from a few nm up to several hundred nm to a spatial precision which is an order of magnitude better than the optical resolution limit; it is *de facto* a super-resolution method. One interesting route for future study could be to use iPAR labelling to explore the effect of chirality on protein assembly processes, for example as is seen in several filamentous biopolymers (137), and even more generally to study “single-molecule cell biology” (138) in the context of “single-molecule cellular biophysics” (139) such as the soft matter properties of cellular material at a single-molecule precise level (140), e.g. stress relaxation effects (104).

There are a range of approaches which have been developed for reporting on aggregate formation in cells. For example, previous studies reporting aggregation of alpha-synuclein amyloid filaments include light-inducible protein clustering system for *in vivo* analysis (141) and multidimensional imaging tools such as and fluorescence lifetime imaging (FLIM) and super-resolution methods such as structured illumination microscopy (SIM) (142) as well as stepwise photobleaching to assess the number of protein subunits present (19). Also, studies involving aggregation effects more generally as stress responses seen in “aggresomes” both in yeast (143) and in bacteria (9, 144).

Although maturation effects of the fluorescent proteins we use here are unlikely to account for more than 10-15% of “dark” fluorescent protein in unstressed cells (59, 88), there are potential physicochemical limitations which may need to be considered. For example, issues with tag folding or localization specifically in high-stress conditions. Similarly, there made be issues relating to spatial variation of pH and molecular crowding, and differences relating to the effects of fluorescent proteins on biomolecular liquid-liquid phase separation (68), which may merit future investigation.

In summary, iPAR offers a robust and improved capability to report on cytoplasmic protein aggregation and shows promising potential to offer new insights into the roles played by stress factors in influencing protein aggregation. We have made the plasmids that encode three fluorescently-tagged variants openly available as a research resource to the scientific community to, we hope, contribute to a wide range of future scientific studies, applicable to a range of advanced fluorescence microscopy modalities (145) including advancing single-molecule biophysics approaches (146, 147) as well as aiding new understanding to the soft matter physics rules behind protein aggregation (71, 140). More generally, our new iPAR technology, has potential to be adapted to other eukaryotic model systems. However, we are careful not to overstate any of the observations we make here in budding yeast in being directly relevant to human cells. Significant additional optimisation is likely to be required to take the iPAR system we have developed here into a human cellular environment if the aim is to directly assess pathology. But with such adaptations there is certainly potential to address several relevant ageing studies and diseases in which protein aggregation is a known or hypothesised factor.

## 5. Ethics

The presented work did not require the use human subject or animals and therefore was not subjected to a welfare committee.

## 6. Data accessibility

Raw data can be openly accessible from Zenodo, DOI: 10.5281/zenodo.10468170 (https://zenodo.org/records/10468171).

Segmentation analysis code can be openly accessible from https://github.com/york-biophysics/ImageJ-Macros (file name: SegSpot.ijm)

The Plasmid construct and cloning maps presented in this article were submitted to Addgene genomic bank under the following ID: 83606 (catalogue number: #212197) and accessible online from https://www.addgene.org/212197/.

## 7. Use of Artificial Intelligence (AI) and AI-assisted technologies

The work presented was not generated using Artificial intelligence tools.

## 8. Authors’ contributions

S.L.: data curation, formal analysis, investigation, methodology, visualization, writing original draft and writing; J.A.L.H.: formal analysis, methodology, visualization and writing; C.M.: funding acquisition, conceptualization, methodology, resources, supervision, visualization and writing; M.C.L.: funding acquisition, conceptualization, methodology, project administration, resources, supervision and writing.

## Acknowledgements

We thank Prof Marija Cvijovic lab and Dr Niek Welkenhuysen for pRS316-*PRC1*-ΔssCPY*-EGFP plasmid donation, Dr Sviatlana Shashkova for donation of the Nrd1-mCherry BY4741 background strain (Chalmers University of Technology, Gothenburg, Sweden) and Maya Schuldiner for the sfGFP-Hof1 BY4741 background strain (Weizmann Institute of Science, Rehovot, Israel). We acknowledge assistance from the Bioscience Technology Facility at the University of York for support with confocal microscopy experiments. We thank Dr Michael Barber (University of York) for assistance with deposition of iPAR plasmids to the Addgene repository.

## Supplementary Information

**Supplementary Figure 1:**
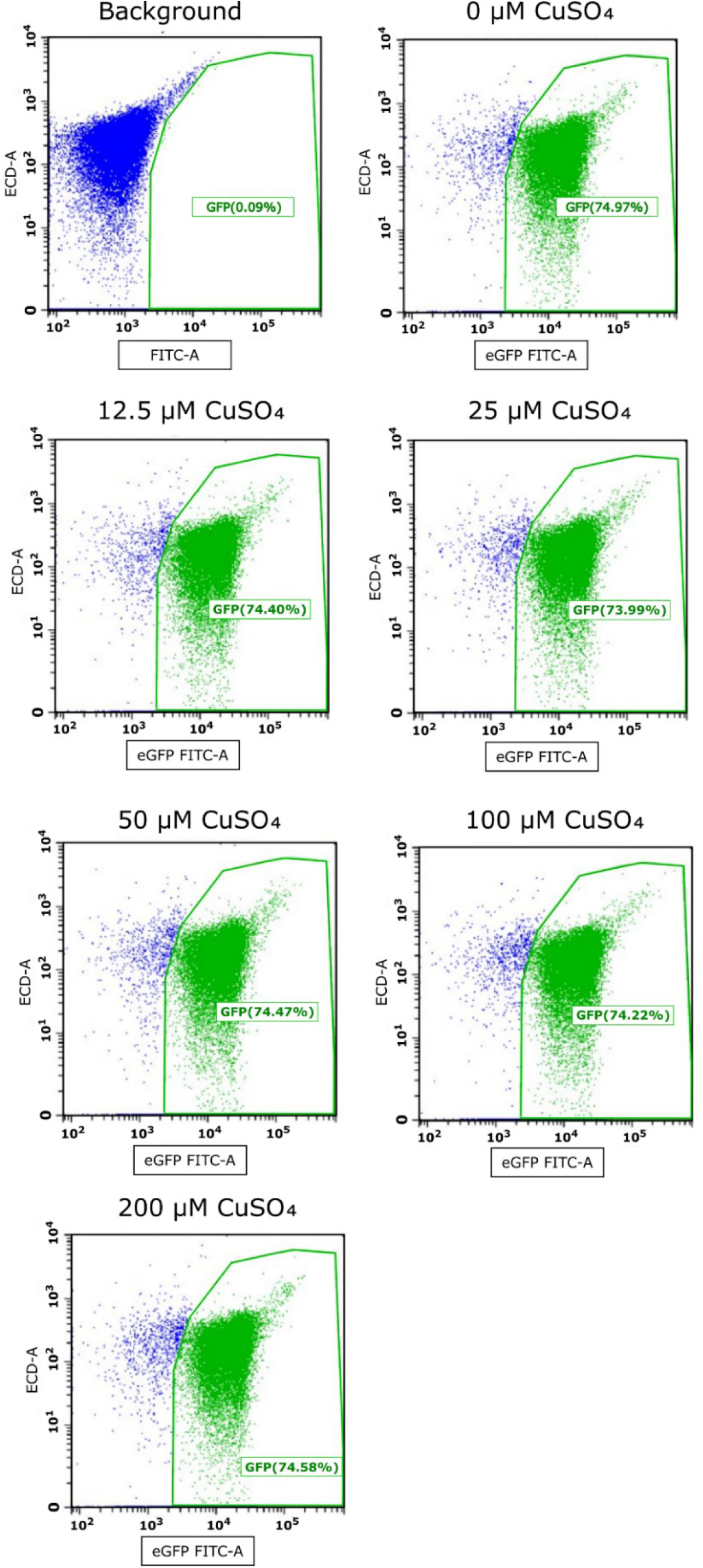
Flow cytometry can quantify fluorescence of cells with high throughput, following induction by copper sulphate. Scatter plot representing the presence of fluorescent positive cells in the cell population analysed by flow cytometry. The background non-fluorescent strain was used to calibrate the presence of non-fluorescent cells, in blue colour. Positive cells expressing Mup1-EGFP are visualised in green and the calculated percentage of EGFP positive is indicated in green.

**Supplementary Figure 2:**
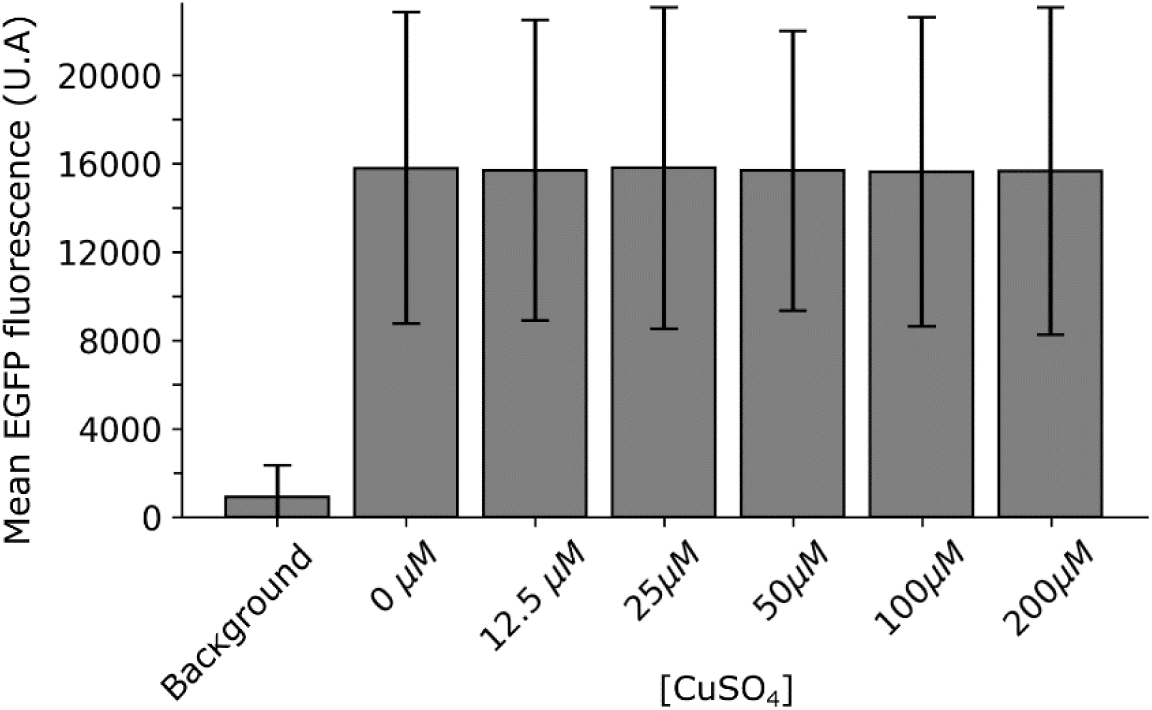
Flow cytometry indicates relative insensitivity to fluorescence brightness for different concentrations of copper sulphate. Box plot representing the mean fluorescence measured in cell population expressing Mup1-EGFP in the presence of different copper sulphate concentrations. Error bar SEM. Number of cells n =>10^4^.

**Supplementary Figure 3:**
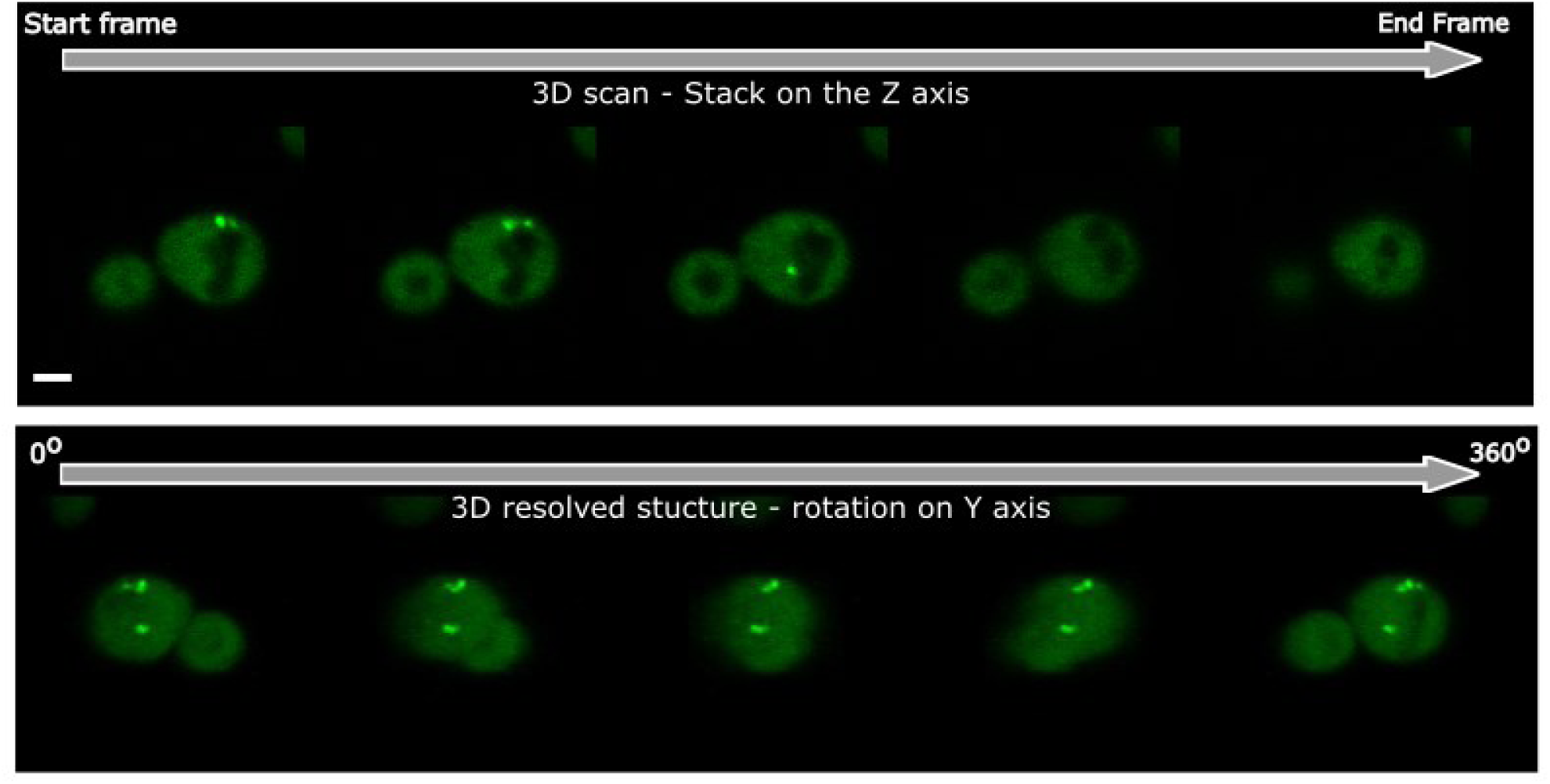
3D visualization can be used to determine the localisation of protein aggregates throughout the full cell volume. Top micrograph: Z stack of protein aggregates for iPAR reporter using mEGFP, 0.33 µm thickness between frames, scales bar: 2 µm. Bottom micrographs: reconstituted 3D volume of the strain, from the Z stack displayed above and using the ImageJ inbuilt 3D project function. See also Supplementary Video 3.

**Supplementary Figure 4:**
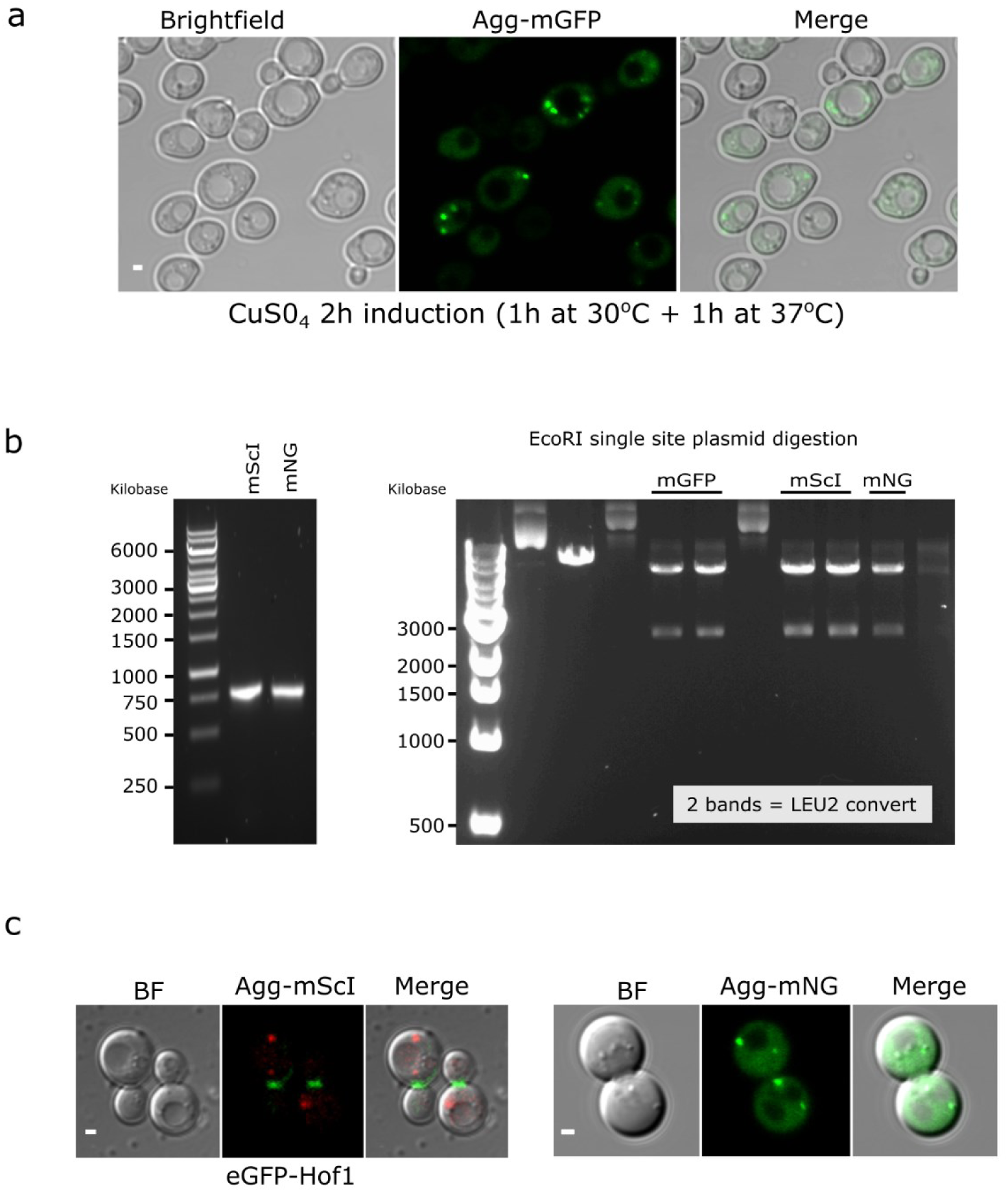
ΔssCPY* aggregate, extension of the strain library, LEU convert and dual colour strain. (a) Micrographs representing selected conditions to induce visible protein aggregates under confocal microscopy imaging. (b) Electrophoresis gels for the construction of fluorescently tagged ΔssCPY* aggregate with either mScarlet-I or mNeonGreen. On the Left: colony PCR verifying fluorophore exchanged after Gibson assembly, mGFP tag sequence being replaced by either mScarlet-I or mNeonGreen (mNG). On the right: Single digest plasmid with EcoRI to verify plasmid Leu conversion, for mGFP, mScarlet-I and mNeonGreen. Plasmid holding the URA selection only cut once and displaying one band. Plasmid holding the LEU selection will instead display two bands. (c) Micrographs mNeonGreen and mScarlet-I version of the ΔssCPY* aggregate reporter. Left: Agg-mScarlet-I expressed in sfGFP-HOF1 strain. Right: Agg-mNeonGreen expressed in BY4742 WT strain.

**Supplementary Figure 5:**
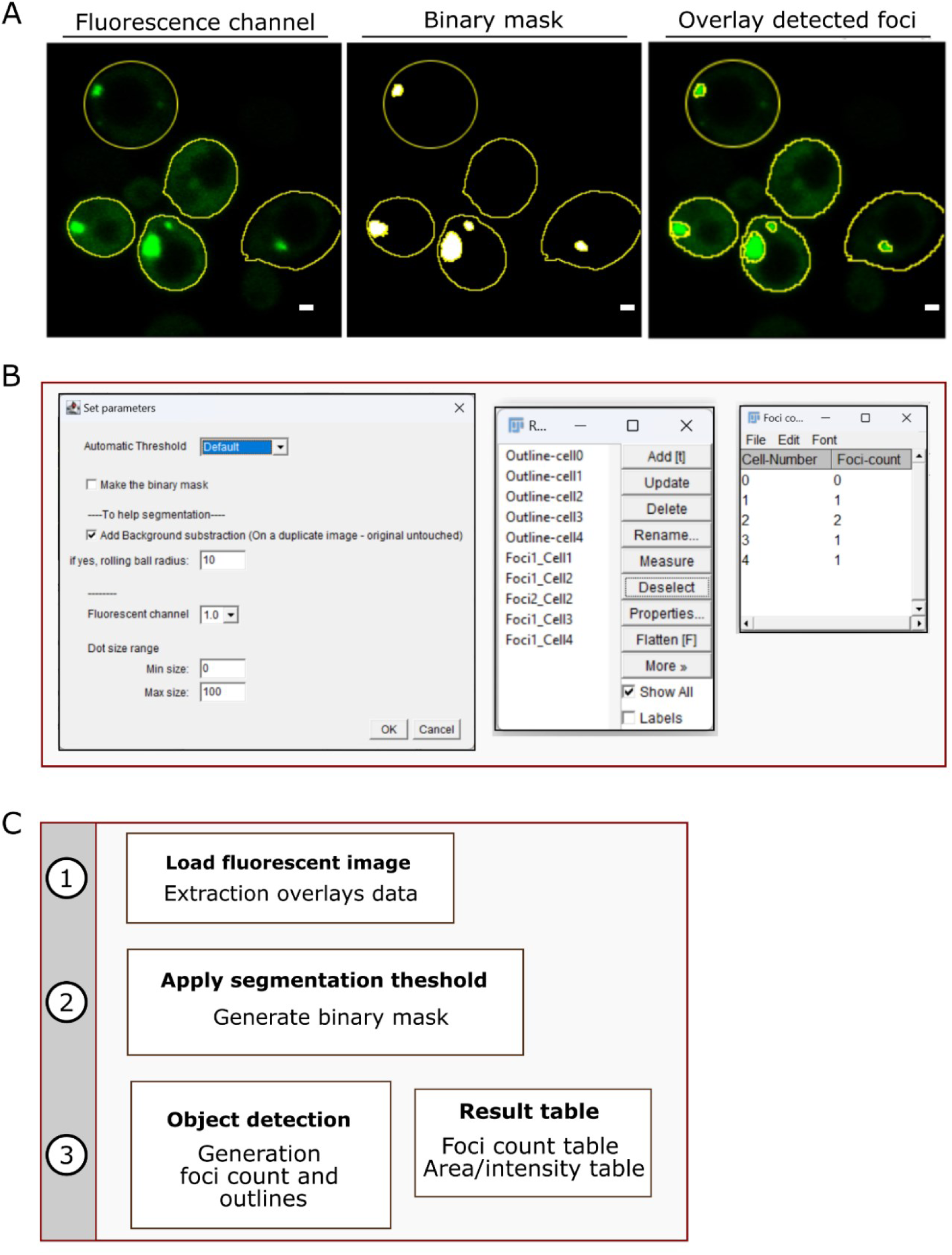
ImageJ macro enables automated, objectivity foci quantification for live cell image data. A) Visual generated segmentation via ImageJ object detection function on the generated binary mask. B: Macro user interface and output sport count table. C) Schematic for the foci detection macro workflow.

**Supplementary Table 1:** Heat shock and aggregates positive cells. Statistical properties including the p-value taken from Student’s *t-*test corresponding to bar plot in Figure3.B

|  | 30°C | 37°C | 42°C |
| --- | --- | --- | --- |
| % aggregates positive cells | 18.98173 | 47.39776 | 58.7738 |
| Standard deviation of % | 5.871798 | 4.943365 | 16.57905 |
| T-test p value (30 vs 37°C) | 7.59 x 10<sup>-5</sup> |  |  |
| T-test p value (37 vs 42°C) |  | 0.261 |  |
| T-test p value (30 vs 42°C) | 5.0 x 10<sup>-3</sup> |  |  |

**Supplementary Table 2:** Heat shock and aggregates counts. Statistical properties including the *t-*test p value corresponding to bar plot in Figure 3.C

|  | 30°C | 37°C | 42°C |
| --- | --- | --- | --- |
| mean foci count/100cells | 26.502 | 62.644 | 131.025 |
| Standard deviation of % | 9.705 | 5.035 | 18.457 |
| T-test p value (30 vs 3°C) | 0.000167 |  |  |
| T-test p value (37 vs 42°C) |  | 0.00021 |  |
| T-test p value (30 vs 42°C) | 2.74 x 10<sup>-5</sup> |  |  |

**Supplementary Table 3:** Comparison aggregates area and intensity properties between mother cells and daughter cells. Statistical properties including the *t-*test p value corresponding to Jitter plots in Figure 5.B

|  | Mother cells | Daughter cells |
| --- | --- | --- |
| Foci mean area | 0.987 | 0.393 |
| Foci median area | 0.822 | 0.284 |
| Standard deviation foci area | 0.744 | 0.291 |
| Foci mean intensity | 52375.69 | 37138.98 |
| Foci median intensity | 51452.68 | 36550.59 |
| Standard deviation foci intensity | 8512.51 | 9785.60 |
| T-test p value (foci area) | 8.60 x 10<sup>-6</sup> |  |
| T-test p value (Foci intensity) | 3 x 10<sup>-14</sup> |  |

**Supplementary Table 4:** Correlation aggregates and key subcellular compartments. Statistical properties including the *t* test p-value corresponding to Jitter plots in Figure 5.D

|  | vacuole | Nucleus |
| --- | --- | --- |
| average correlation percentage (%) | 43.94737 | 68.11404 |
| Standard deviation | 36.34713 | 32.67815 |
| T-test p value | 0.0057 |  |

## References

1. Amm I, Sommer T, Wolf DH. Protein quality control and elimination of protein waste: The role of the ubiquitin–proteasome system. Biochimica et Biophysica Acta (BBA)-Molecular Cell Research. 2014;1843(1):182–96.

2. Bukau B, Weissman J, Horwich A. Molecular chaperones and protein quality control. Cell. 2006;125(3):443–51.

3. Chen B, Retzlaff M, Roos T, Frydman J. Cellular strategies of protein quality control. Cold Spring Harbor Perspectives in Biology. 2011;3(8):a004374.

4. Karmon O, Ben Aroya S. Spatial organization of proteasome aggregates in the regulation of proteasome homeostasis. Frontiers in Molecular Biosciences. 2020;6:150.

5. Kwon YT, Ciechanover A. The ubiquitin code in the ubiquitin-proteasome system and autophagy. Trends in Biochemical Sciences. 2017;42(11):873–86.

6. Mizushima N, Komatsu M. Autophagy: renovation of cells and tissues. Cell. 2011;147(4):728–41.

7. Fink AL. Protein aggregation: folding aggregates, inclusion bodies and amyloid. Folding and Design. 1998;3(1):R9–R23.

8. Sluzky V, Tamada JA, Klibanov AM, Langer R. Kinetics of insulin aggregation in aqueous solutions upon agitation in the presence of hydrophobic surfaces. Proceedings of the National Academy of Sciences. 1991;88(21):9377–81.

9. Jin X, Lee J-E, Schaefer C, Luo X, Wollman AJ, Payne-Dwyer AL, et al. Membraneless organelles formed by liquid-liquid phase separation increase bacterial fitness. Science Advances. 2021;7(43):eabh2929.

10. Lee S, Choi MC, Al Adem K, Lukman S, Kim T-Y. Aggregation and cellular toxicity of pathogenic or non-pathogenic proteins. Scientific Reports. 2020;10(1):5120.

11. Holmes WM, Klaips CL, Serio TR. Defining the limits: Protein aggregation and toxicity in vivo. Critical Reviews in Biochemistry and Molecular Biology. 2014;49(4):294–303.

12. Hipp MS, Park S-H, Hartl FU. Proteostasis impairment in protein-misfolding and-aggregation diseases. Trends in Cell Biology. 2014;24(9):506–14.

13. Hartl FU. Protein Misfolding Diseases. Annual Review of Biochemistry. 2017;86:21–6.

14. Dobson CM. The structural basis of protein folding and its links with human disease. Philosophical Transactions of the Royal Society of London Series B: Biological Sciences. 2001;356(1406):133–45.

15. Dobson CM. Protein misfolding, evolution and disease. Trends in Biochemical Sciences. 1999;24(9):329–32.

16. Ross CA, Poirier MA. Protein aggregation and neurodegenerative disease. Nature Medicine. 2004;10(Suppl 7):S10–S7.

17. Chiti F, Dobson CM. Protein misfolding, functional amyloid, and human disease. Annual Review of Biochemistry. 2006;75:333–66.

18. Chiti F, Dobson CM. Protein misfolding, amyloid formation, and human disease: a summary of progress over the last decade. Annual Review of Biochemistry. 2017;86:27–68.

19. Dresser L, Hunter P, Yendybayeva F, Hargreaves AL, Howard JA, Evans GJ, et al. Amyloid-β oligomerization monitored by single-molecule stepwise photobleaching. Methods. 2021;193:80–95.

20. Budnar P, Tangirala R, Bakthisaran R, Rao CM. Protein aggregation and cataract: Role of age-related modifications and mutations in α-crystallins. Biochemistry (Moscow). 2022;87(3):225–41.

21. Arrasate M, Finkbeiner S. Protein aggregates in Huntington’s disease. Experimental Neurology. 2012;238(1):1–11.

22. Moreau KL, King JA. Protein misfolding and aggregation in cataract disease and prospects for prevention. Trends in Molecular Medicine. 2012;18(5):273–82.

23. Surguchev A, Surguchov A. Conformational diseases: looking into the eyes. Brain Research Bulletin. 2010;81(1):12–24.

24. Zimmermann A, Hofer S, Pendl T, Kainz K, Madeo F, Carmona-Gutierrez D. Yeast as a tool to identify anti-aging compounds. FEMS Yeast Research. 2018;18(6):foy020.

25. Perocchi F, Mancera E, Steinmetz LM. Systematic screens for human disease genes, from yeast to human and back. Molecular bioSystems. 2008;4(1):18–29.

26. Schmitt ME, Clayton DA. Conserved features of yeast and mammalian mitochondrial DNA replication. Current Opinion in Genetics & Development. 1993;3(5):769–74.

27. Boos D, Sanchez-Pulido L, Rappas M, Pearl LH, Oliver AW, Ponting CP, Diffley JF. Regulation of DNA replication through Sld3-Dpb11 interaction is conserved from yeast to humans. Current Biology. 2011;21(13):1152–7.

28. Goodman AJ, Daugharthy ER, Kim J. Pervasive antisense transcription is evolutionarily conserved in budding yeast. Molecular Biology and Evolution. 2013;30(2):409–21.

29. Hahn S, Young ET. Transcriptional regulation in Saccharomyces cerevisiae: transcription factor regulation and function, mechanisms of initiation, and roles of activators and coactivators. Genetics. 2011;189(3):705–36.

30. Bennett MK, Scheller RH. The molecular machinery for secretion is conserved from yeast to neurons. Proceedings of the National Academy of Sciences. 1993;90(7):2559–63.

31. Ma M, Burd CG. Retrograde trafficking and plasma membrane recycling pathways of the budding yeast Saccharomyces cerevisiae. Traffic. 2020;21(1):45–59.

32. MacDonald C, Piper RC. Genetic dissection of early endosomal recycling highlights a TORC1-independent role for Rag GTPases. Journal of Cell Biology. 2017;216(10):3275–90.

33. Delic M, Valli M, Graf AB, Pfeffer M, Mattanovich D, Gasser B. The secretory pathway: exploring yeast diversity. FEMS Microbiology Reviews. 2013;37(6):872–914.

34. Benyair R, Ron E, Lederkremer GZ. Protein quality control, retention, and degradation at the endoplasmic reticulum. International Review of Cell and Molecular Biology. 2011;292:197–280.

35. Tyedmers J, Mogk A, Bukau B. Cellular strategies for controlling protein aggregation. Nature Reviews Molecular Cell Biology. 2010;11(11):777–88.

36. Chakrabarti A, Chen AW, Varner JD. A review of the mammalian unfolded protein response. Biotechnology and Bioengineering. 2011;108(12):2777–93.

37. Fink AL. Chaperone-mediated protein folding. Physiological Reviews. 1999;79(2):425–49.

38. Higgins R, Kabbaj M-H, Hatcher A, Wang Y. The absence of specific yeast heat-shock proteins leads to abnormal aggregation and compromised autophagic clearance of mutant Huntingtin proteins. PLOS ONE. 2018;13(1):e0191490.

39. Muchowski PJ, Schaffar G, Sittler A, Wanker EE, Hayer-Hartl MK, Hartl FU. Hsp70 and hsp40 chaperones can inhibit self-assembly of polyglutamine proteins into amyloid-like fibrils. Proceedings of the National Academy of Sciences. 2000;97(14):7841–6.

40. Park S-H, Bolender N, Eisele F, Kostova Z, Takeuchi J, Coffino P, Wolf DH. The cytoplasmic Hsp70 chaperone machinery subjects misfolded and endoplasmic reticulum import-incompetent proteins to degradation via the ubiquitin–proteasome system. Molecular Biology of the Cell. 2007;18(1):153–65.

41. Spence J, Sadis S, Haas AL, Finley D. A ubiquitin mutant with specific defects in DNA repair and multiubiquitination. Molecular and Cellular Biology. 1995;15(3):1265–73.

42. McClellan AJ, Tam S, Kaganovich D, Frydman J. Protein quality control: chaperones culling corrupt conformations. Nature Cell Biology. 2005;7(8):736–41.

43. Schneider KL, Wollman AJ, Nyström T, Shashkova S. Comparison of endogenously expressed fluorescent protein fusions behaviour for protein quality control and cellular ageing research. Scientific Reports. 2021;11(1):12819.

44. Stolz A, Wolf DH. Use of CPY* and its derivatives to study protein quality control in various cell compartments. Ubiquitin Family Modifiers and the Proteasome: Reviews and Protocols. 2012:489–504.

45. Bowman S, Churcher C, Badcock K, Brown D, Chillingworth T, Connor R, et al. The nucleotide sequence of Saccharomyces cerevisiae chromosome XIII. Nature. 1997;387(6632 Suppl):90–3.

46. Valls LA, Hunter CP, Rothman JH, Stevens TH. Protein sorting in yeast: the localization determinant of yeast vacuolar carboxypeptidase Y resides in the propeptide. Cell. 1987;48(5):887–97.

47. Finger A, Knop M, Wolf DH. Analysis of two mutated vacuolar proteins reveals a degradation pathway in the endoplasmic reticulum or a related compartment of yeast. European Journal of Biochemistry. 1993;218(2):565–74.

48. Medicherla B, Kostova Z, Schaefer A, Wolf DH. A genomic screen identifies Dsk2p and Rad23p as essential components of ER-associated degradation. EMBO Reports. 2004;5(7):692–7.

49. Park S-H. Molecular chaperones in protein quality control: from recognition to degradation: PhD Thesis: University of Stuttgart; 2007.

50. Eisele F. Components and mechanisms of cytoplasmic protein quality control and elimination of regulatory enzymes: PhD Thesis: University of Stuttgart; 2011.

51. Öling D, Eisele F, Kvint K, Nyström T. Opposing roles of U bp3-dependent deubiquitination regulate replicative life span and heat resistance. The EMBO Journal. 2014;33(7):747–61.

52. Hanzén S. Proteostasis and Aging in Saccharomyces cerevisiae-The role of a Peroxiredoxin: PhD Thesis: University of Gothenburg; 2017.

53. Schnitzer B, Welkenhuysen N, Leake MC, Shashkova S, Cvijovic M. The effect of stress on biophysical characteristics of misfolded protein aggregates in living Saccharomyces cerevisiae cells. Experimental Gerontology. 2022;162:111755.

54. Bradley PH, Brauer MJ, Rabinowitz JD, Troyanskaya OG. Coordinated concentration changes of transcripts and metabolites in Saccharomyces cerevisiae. PLoS Computational Biology. 2009;5(1):e1000270.

55. Segal E, Shapira M, Regev A, Pe’er D, Botstein D, Koller D, Friedman N. Module networks: identifying regulatory modules and their condition-specific regulators from gene expression data. Nature Genetics. 2003;34(2):166–76.

56. Costantini LM, Fossati M, Francolini M, Snapp EL. Assessing the tendency of fluorescent proteins to oligomerize under physiologic conditions. Traffic. 2012;13(5):643–9.

57. Cranfill PJ, Sell BR, Baird MA, Allen JR, Lavagnino Z, De Gruiter HM, et al. Quantitative assessment of fluorescent proteins. Nature methods. 2016;13(7):557–62.

58. Pope JR, Johnson RL, Jamieson WD, Worthy HL, Kailasam S, Ahmed RD, et al. Association of fluorescent protein pairs and its significant impact on fluorescence and energy transfer. Advanced Science. 2021;8(1):2003167.

59. Shashkova S, Wollman AJ, Hohmann S, Leake MC. Characterising Maturation of GFP and mCherry of Genomically Integrated Fusions in Saccharomyces cerevisiae. Bio-Protocol. 2018;8(2):e2710.

60. Reyes-Lamothe R, Sherratt DJ, Leake MC. Stoichiometry and architecture of active DNA replication machinery in *Escherichia coli*. Science. 2010;328(5977):498–501.

61. Stracy M, Adam J, Kaja E, Gapinski J, Lee J-E, Leek VA, et al. Single-molecule imaging of DNA gyrase activity in living *Escherichia coli*. Nucleic Acids Research. 2019;47(1):210–20.

62. Badrinarayanan A, Reyes-Lamothe R, Uphoff S, Leake MC, Sherratt DJ. *In vivo* architecture and action of bacterial structural maintenance of chromosome proteins. Science. 2012;338(6106):528–31.

63. Sun Y, Wollman AJM, Huang F, Leake MC, Liu L-N. Single-Organelle Quantification Reveals Stoichiometric and Structural Variability of Carboxysomes Dependent on the Environment. The Plant Cell. 2019;31(7):1648–64.

64. Payne-Dwyer AL, Syeda AH, Shepherd JW, Frame L, Leake MC. RecA and RecB: probing complexes of DNA repair proteins with mitomycin C in live *Escherichia coli* with single-molecule sensitivity. Journal of The Royal Society Interface. 2022;19(193).

65. Wollman AJM, Muchová K, Chromiková Z, Wilkinson AJ, Barák I, Leake MC. Single-molecule optical microscopy of protein dynamics and computational analysis of images to determine cell structure development in differentiating *Bacillus subtilis*. Computational and Structural Biotechnology Journal. 2020;18:1474–86.

66. Syeda AH, Wollman AJM, Hargreaves AL, Howard JAL, Brüning J-G, McGlynn P, Leake MC. Single-molecule live cell imaging of Rep reveals the dynamic interplay between an accessory replicative helicase and the replisome. Nucleic Acids Research. 2019;47(12):6287–98.

67. Wollman AJM, Syeda AH, Howard JAL, Payne-Dwyer A, Leech A, Warecka D, et al. Tetrameric UvrD Helicase Is Located at the E. Coli Replisome due to Frequent Replication Blocks. Journal of Molecular Biology. 2024;436(2):168369.

68. Leake MC. Transcription factors in eukaryotic cells can functionally regulate gene expression by acting in oligomeric assemblies formed from an intrinsically disordered protein phase transition enabled by molecular crowding. Transcr. 2018;9(5):298–306.

69. Shashkova S, Nyström T, Leake MC, Wollman AJ. Correlative single-molecule fluorescence barcoding of gene regulation in Saccharomyces cerevisiae. Methods. 2021;193:62–7.

70. Shashkova S, Wollman AJM, Leake MC, Hohmann S. The yeast Mig1 transcriptional repressor is dephosphorylated by glucose-dependent and -independent mechanisms. FEMS Microbiology Letters. 2017;364(14).

71. Payne-Dwyer A, Kumar G, Barrett J, Gherman LK, Hodgkinson M, Plevin M, et al. Predicting Rubisco-Linker Condensation from Titration in the Dilute Phase. Physical Review Letters. 2024;132(21).

72. Adler L, Lau CS, Shaikh KM, Maldegem KAv, Payne-Dwyer AL, Lefoulon C, et al. The role of BST4 in the pyrenoid of *Chlamydomonas reinhardtii*. Plant Physiology. 2024;In Press.

73. Wollman AJM, Fournier C, Llorente-Garcia I, Harriman O, Payne-Dwyer AL, Shashkova S, et al. Critical roles for EGFR and EGFR–HER2 clusters in EGF binding of SW620 human carcinoma cells. Journal of The Royal Society Interface. 2022;19(190).

74. Hunter P, Payne-Dwyer AL, Shaw M, Signoret N, Leake MC. Single-molecule and super-resolved imaging deciphers membrane behavior of onco-immunogenic CCR5. iScience. 2022;25(12):105675.

75. Cosgrove J, Novkovic M, Albrecht S, Pikor NB, Zhou Z, Onder L, et al. B cell zone reticular cell microenvironments shape CXCL13 gradient formation. Nature communications. 2020;11(1).

76. Payne-Dwyer AL, Jang G-J, Dean C, Leake MC. SlimVar: rapid *in vivo* single-molecule tracking of chromatin regulators in plants. bioRxiv. 2024:10.1101/2024.05.17.594710.

77. Miller H, Cosgrove J, Wollman AJM, Taylor E, Zhou Z, O’Toole PJ, et al. High-Speed Single-Molecule Tracking of CXCL13 in the B-Follicle. Frontiers in Immunology. 2018;9.

78. Brachmann CB, Davies A, Cost GJ, Caputo E, Li J, Hieter P, Boeke JD. Designer deletion strains derived from Saccharomyces cerevisiae S288C: a useful set of strains and plasmids for PCR-mediated gene disruption and other applications. Yeast. 1998;14(2):115–32.

79. Weill U, Yofe I, Sass E, Stynen B, Davidi D, Natarajan J, et al. Genome-wide SWAp-Tag yeast libraries for proteome exploration. Nature Methods. 2018;15(8):617–22.

80. Laidlaw KM, Bisinski DD, Shashkova S, Paine KM, Veillon MA, Leake MC, MacDonald C. A glucose-starvation response governs endocytic trafficking and eisosomal retention of surface cargoes in budding yeast. Journal of Cell Science. 2021;134(2):jcs257733.

81. MacDonald C, Payne JA, Aboian M, Smith W, Katzmann DJ, Piper RC. A family of tetraspans organizes cargo for sorting into multivesicular bodies. Developmental Cell. 2015;33(3):328–42.

82. Landgraf D, Okumus B, Chien P, Baker TA, Paulsson J. Segregation of molecules at cell division reveals native protein localization. Nature Methods. 2012;9(5):480–2.

83. Gibson DG, Young L, Chuang R-Y, Venter JC, Hutchison III CA, Smith HO. Enzymatic assembly of DNA molecules up to several hundred kilobases. Nature Methods. 2009;6(5):343–5.

84. Bindels DS, Haarbosch L, Van Weeren L, Postma M, Wiese KE, Mastop M, et al. mScarlet: a bright monomeric red fluorescent protein for cellular imaging. Nature Methods. 2017;14(1):53–6.

85. Shaner NC, Lambert GG, Chammas A, Ni Y, Cranfill PJ, Baird MA, et al. A bright monomeric green fluorescent protein derived from Branchiostoma lanceolatum. Nature Methods. 2013;10(5):407–9.

86. Fogel S, Welch JW. Tandem gene amplification mediates copper resistance in yeast. Proceedings of the National Academy of Sciences. 1982;79(17):5342–6.

87. Vida TA, Emr SD. A new vital stain for visualizing vacuolar membrane dynamics and endocytosis in yeast. The Journal of Cell Biology. 1995;128(5):779–92.

88. Leake MC, Chandler JH, Wadhams GH, Bai F, Berry RM, Armitage JP. Stoichiometry and turnover in single, functioning membrane protein complexes. Nature. 2006;443(7109):355–8.

89. Wollman AJM, Hedlund EG, Shashkova S, Leake MC. Towards mapping the 3D genome through high speed single-molecule tracking of functional transcription factors in single living cells. Methods. 2020;170:82–9.

90. Shepherd JW, Lecinski S, Wragg J, Shashkova S, MacDonald C, Leake MC. Molecular crowding in single eukaryotic cells: Using cell environment biosensing and single-molecule optical microscopy to probe dependence on extracellular ionic strength, local glucose conditions, and sensor copy number. Methods. 2021;193:54–61.

91. Laidlaw KME, Calder G, MacDonald C. Recycling of cell surface membrane proteins from yeast endosomes is regulated by ubiquitinated Ist1. Journal of Cell Biology. 2022;221(11).

92. Lecinski S, Shepherd JW, Frame L, Hayton I, MacDonald C, Leake MC. Investigating molecular crowding during cell division and hyperosmotic stress in budding yeast with FRET. Current Topics in Membranes. 88: Elsevier; 2021. p. 75–118.

93. Walker T. Cell Magic Wand. https://github.com/fitzlab/CellMagicWand2016 [

94. Plank M, Wadhams GH, Leake MC. Millisecond timescale slimfield imaging and automated quantification of single fluorescent protein molecules for use in probing complex biological processes. Integrative Biology. 2009;1(10):602–12.

95. Wollman AJ, Shashkova S, Hedlund EG, Friemann R, Hohmann S, Leake MC. Transcription factor clusters regulate genes in eukaryotic cells. Elife. 2019;8:e45804.

96. Wollman AJM, Leake MC. Single-Molecule Narrow-Field Microscopy of Protein-DNA Binding Dynamics in Glucose Signal Transduction of Live Yeast Cells. Methods in Molecular Biology: Springer US; 2022. p. 5–16.

97. Miller H, Zhou Z, Wollman AJ, Leake MC. Superresolution imaging of single DNA molecules using stochastic photoblinking of minor groove and intercalating dyes. Methods. 2015;88:81–8.

98. Shepherd JW, Higgins EJ, Wollman AJM, Leake MC. PySTACHIO: Python Single-molecule TrAcking stoiCHiometry Intensity and simulatiOn, a flexible, extensible, beginner-friendly and optimized program for analysis of single-molecule microscopy data. Computational and Structural Biotechnology Journal. 2021;19:4049–58.

99. Sowa Y, Rowe AD, Leake MC, Yakushi T, Homma M, Ishijima A, Berry RM. Direct observation of steps in rotation of the bacterial flagellar motor. Nature. 2005;437(7060):916–9.

100. Wollman AJM, Leake MC. Millisecond single-molecule localization microscopy combined with convolution analysis and automated image segmentation to determine protein concentrations in complexly structured, functional cells, one cell at a time. Faraday Discussions. 2015;184:401–24.

101. Leake MC. Analytical tools for single-molecule fluorescence imaging *in cellulo*. Physical Chemistry Chemical Physics. 2014;16(25):12635–47.

102. Llorente-Garcia I, Lenn T, Erhardt H, Harriman OL, Liu LN, Robson A, et al. Single-molecule *in vivo* imaging of bacterial respiratory complexes indicates delocalized oxidative phosphorylation. Biochimica et biophysica acta. 2014;1837(6):811–24.

103. Leake MC, Greene NP, Godun RM, Granjon T, Buchanan G, Chen S, et al. Variable stoichiometry of the TatA component of the twin-arginine protein transport system observed by *in vivo* single-molecule imaging. Proceedings of the National Academy of Sciences. 2008;105(40):15376–81.

104. Linke WA, Leake MC. Multiple sources of passive stress relaxation in muscle fibres. Physics in Medicine & Biology. 2004;49(16):3613–27.

105. Robinson JS, Klionsky DJ, Banta LM, Emr SD. Protein sorting in Saccharomyces cerevisiae: isolation of mutants defective in the delivery and processing of multiple vacuolar hydrolases. Molecular and Cellular Biology. 1988;8(11):4936–48.

106. Rothman JH, Stevens TH. Protein sorting in yeast: mutants defective in vacuole biogenesis mislocalize vacuolar proteins into the late secretory pathway. Cell. 1986;47(6):1041–51.

107. Rothman JE. Polypeptide chain binding proteins: catalysts of protein folding and related processes in cells. Cell. 1989;59(4):591–601.

108. Bankaitis VA, Johnson LM, Emr SD. Isolation of yeast mutants defective in protein targeting to the vacuole. Proceedings of the National Academy of Sciences. 1986;83(23):9075–9.

109. Stevens T, Esmon B, Schekman R. Early stages in the yeast secretory pathway are required for transport of carboxypeptidase Y to the vacuole. Cell. 1982;30(2):439–48.

110. Paxman J, Zhou Z, O’Laughlin R, Liu Y, Li Y, Tian W, et al. Age-dependent aggregation of ribosomal RNA-binding proteins links deterioration in chromatin stability with challenges to proteostasis. eLife. 2022;11:e75978.

111. Schnitzer B, Welkenhuysen N, Leake MC, Shashkova S, Cvijovic M. The effect of stress on biophysical characteristics of misfolded protein aggregates in living Saccharomyces cerevisiae cells. Experimental gerontology. 2022:111755.

112. Valls LA, Hunter CP, Rothman JH, Stevens TH. Protein sorting in yeast: the localization determinant of yeast vacuolar carboxypeptidase Y resides in the propeptide. Cell. 1987;48(5):887–97.

113. Macreadie IG, Horaitis O, Vaughan PR, Des Clark-Walker G. Constitutive expression of the Saccharomyces cerevisiae CUP1 gene in Kluyveromyces lactis. Yeast. 1991;7(2):127–35.

114. MacDonald C, Winistorfer S, Pope RM, Wright ME, Piper RC. Enzyme reversal to explore the function of yeast E3 ubiquitin-ligases. Traffic. 2017;18(7):465–84.

115. Reith P, Braam S, Welkenhuysen N, Lecinski S, Shepherd J, MacDonald C, et al. The effect of lithium on the budding yeast Saccharomyces cerevisiae upon stress adaptation. Microorganisms. 2022;10(3):590.

116. Vinutha H, Poornima B, Sagar B, editors. Detection of outliers using interquartile range technique from intrusion dataset. Information and Decision Sciences: Proceedings of the 6th International Conference on FICTA; 2018: Springer.

117. Schneider KL, Nyström T, Widlund PO. Studying spatial protein quality control, proteopathies, and aging using different model misfolding proteins in S. cerevisiae. Frontiers in Molecular Neuroscience. 2018;11:249.

118. Spokoini R, Moldavski O, Nahmias Y, England JL, Schuldiner M, Kaganovich D. Confinement to organelle-associated inclusion structures mediates asymmetric inheritance of aggregated protein in budding yeast. Cell Reports. 2012;2(4):738–47.

119. Hill SM, Hanzén S, Nyström T. Restricted access: spatial sequestration of damaged proteins during stress and aging. EMBO Reports. 2017;18(3):377–91.

120. Sontag EM, Samant RS, Frydman J. Mechanisms and functions of spatial protein quality control. Annual Review of Biochemistry. 2017;86:97–122.

121. Rothe S, Prakash A, Tyedmers J. The insoluble protein deposit (IPOD) in yeast. Frontiers in Molecular Neuroscience. 2018:237.

122. Wollman AJM, Shashkova S, Hedlund EG, Friemann R, Hohmann S, Leake MC. Transcription factor clusters regulate genes in eukaryotic cells. eLife. 2019;8:e45804.

123. Li KW, Lu MS, Iwamoto Y, Drubin DG, Pedersen RT. A preferred sequence for organelle inheritance during polarized cell growth. Journal of Cell Science. 2021;134(21):jcs258856.

124. Clay L, Caudron F, Denoth-Lippuner A, Boettcher B, Buvelot Frei S, Snapp EL, Barral Y. A sphingolipid-dependent diffusion barrier confines ER stress to the yeast mother cell. elife. 2014;3:e01883.

125. Ouellet J, Barral Y. Organelle segregation during mitosis: lessons from asymmetrically dividing cells. Journal of Cell Biology. 2012;196(3):305–13.

126. Nyström T. Spatial protein quality control and the evolution of lineage-specific ageing. Philosophical Transactions of the Royal Society B: Biological Sciences. 2011;366(1561):71–5.

127. Nyström T, Liu B. The mystery of aging and rejuvenation—a budding topic. Current Opinion in Microbiology. 2014;18:61–7.

128. Labbadia J, Morimoto RI. The biology of proteostasis in aging and disease. Annual Review of Biochemistry. 2015;84:435–64.

129. Balchin D, Hayer-Hartl M, Hartl FU. In vivo aspects of protein folding and quality control. Science. 2016;353(6294):aac4354.

130. Sontag EM, Vonk WI, Frydman J. Sorting out the trash: the spatial nature of eukaryotic protein quality control. Current Opinion in Cell Biology. 2014;26:139–46.

131. Hohmann S. Osmotic stress signaling and osmoadaptation in yeasts. Microbiol Mol Biol Rev. 2002;66(2):300–72.

132. Shashkova S, Andersson M, Hohmann S, Leake MC. Correlating single-molecule characteristics of the yeast aquaglyceroporin Fps1 with environmental perturbations directly in living cells. Methods. 2021;193:46–53.

133. Ogrodnik M, Salmonowicz H, Brown R, Turkowska J, Średniawa W, Pattabiraman S, et al. Dynamic JUNQ inclusion bodies are asymmetrically inherited in mammalian cell lines through the asymmetric partitioning of vimentin. Proceedings of the National Academy of Sciences. 2014;111(22):8049–54.

134. Lecinski S, Shepherd JW, Bunting K, Dresser L, Quinn SD, MacDonald C, Leake MC. Correlating viscosity and molecular crowding with fluorescent nanobeads and molecular probes: in vitro and in vivo. Interface Focus. 2022;12(6):20220042.

135. Leake MC. Biophysics Tools and Techniques for the Physics of Life (Second Edition): CRC Press; 2023 12/12/2023. 456 p.

136. Shashkova S, Leake MC. Single-molecule fluorescence microscopy review: shedding new light on old problems. Biosci Rep. 2017;37(4).

137. Shepherd JW, Guilbaud S, Zhou Z, Howard JAL, Burman M, Schaefer C, et al. Correlating fluorescence microscopy, optical and magnetic tweezers to study single chiral biopolymers such as DNA. Nat Commun. 2024;15(1):2748.

138. Leake MC. Shining the spotlight on functional molecular complexes: The new science of single-molecule cell biology. Commun Integr Biol. 2010;3(5):415–8.

139. Leake MC. Single-Molecule Cellular Biophysics. In: Leake MC, editor. Single-Molecule Cellular Biophysics. Cambridge: Cambridge University Press; 2013. p. i-vi.

140. Harriman OL, Leake MC. Single molecule experimentation in biological physics: exploring the living component of soft condensed matter one molecule at a time. Journal of Physics: Condensed Matter. 2011;23(50):503101.

141. Bérard M, Sheta R, Malvaut S, Rodriguez-Aller R, Teixeira M, Idi W, et al. A light-inducible protein clustering system for in vivo analysis of α-synuclein aggregation in Parkinson disease. PLoS biology. 2022;20(3):e3001578.

142. Lu M, Williamson N, Mishra A, Michel CH, Kaminski CF, Tunnacliffe A, Kaminski Schierle GS. Structural progression of amyloid-β Arctic mutant aggregation in cells revealed by multiparametric imaging. J Biol Chem. 2019;294(5):1478–87.

143. Lu M, Boschetti C, Tunnacliffe A. Long Term Aggresome Accumulation Leads to DNA Damage, p53-dependent Cell Cycle Arrest, and Steric Interference in Mitosis. J Biol Chem. 2015;290(46):27986–8000.

144. Pu Y, Li Y, Jin X, Tian T, Ma Q, Zhao Z, et al. ATP-Dependent Dynamic Protein Aggregation Regulates Bacterial Dormancy Depth Critical for Antibiotic Tolerance. Mol Cell. 2019;73(1):143–56.e4.

145. Bose KS, Sarma RH. Delineation of the intimate details of the backbone conformation of pyridine nucleotide coenzymes in aqueous solution. Biochem Biophys Res Commun. 1975;66(4):1173–9.

146. Miller H, Zhou Z, Shepherd J, Wollman AJM, Leake MC. Single-molecule techniques in biophysics: a review of the progress in methods and applications. Reports on Progress in Physics. 2018;81(2):024601.

147. Leake MC. The physics of life: one molecule at a time. Philosophical Transactions of the Royal Society B: Biological Sciences. 2013;368(1611):20120248.

